# Host-parasite coevolution promotes innovation through deformations in fitness landscapes

**DOI:** 10.1101/2021.06.25.449783

**Authors:** Animesh Gupta, Luis Zaman, Hannah M. Strobel, Jenna Gallie, Alita R. Burmeister, Benjamin Kerr, Einat S. Tamar, Roy Kishony, Justin R. Meyer

## Abstract

During the struggle for survival, populations occasionally evolve new functions that give them access to untapped ecological opportunities. Theory suggests that coevolution between species can promote the evolution of such innovations by deforming fitness landscapes in ways that open new adaptive pathways. We directly tested this idea by using high throughput gene editing-phenotyping technology (MAGE-Seq) to measure the fitness landscape of a virus, bacteriophage λ, as it coevolved with its host, the bacterium *Escherichia coli*. Through computer simulations of λ’s evolution on the empirical fitness landscape, we showed that λ was more likely to evolve to use a new receptor if it experienced a shift in its fitness landscape caused by coevolution. This result was further validated by additional laboratory experiments. This study provides direct evidence for the role of coevolution in driving evolutionary novelty and provides a quantitative framework for predicting evolution in coevolving ecological communities.

## Main Text

A starting point for understanding how populations evolve is to assume that they exist in an unchanging world where they can adapt towards optimality^1,2^. However, even in static environments, populations never reach optimality because their circumstances continuously change as neighbouring species coevolve with them^3^. This more dynamic view of the evolutionary process opens up the potential for unbounded evolution and creates new opportunities for evolutionary innovation^4–8^. Darwin recognized this potential in the final pages of *On the Origin of Species* where he wrote that, “It is interesting to contemplate an entangled bank” of organisms evolving with one another to produce such a variety of forms and functions^9^. But he also realized the empirical challenges created by the richness of species interactions within ecological communities in his further description of “these elaborately constructed forms,… dependent on each other in so complex a manner…”^9^. The complexity arises because an organism’s fitness is a function of its interactions with other species, and the strength and form of these interactions continuously change as they coevolve. Furthermore, the coevolving traits of the organisms are encoded within genomes by mutations that might interact with one another, a pervasive phenomenon called epistasis^10^. This means that interactions at all levels must be considered; from mutation-by-mutation within a species (classical epistasis), to mutation-by-mutation between species (interspecific epistasis), and higher order phenomena such as classic and interspecific epistasis combined.

Many advances have been made over recent decades that enable us to tackle this combinatorial problem. Efficient genetic engineering methods permit the construction of genetic libraries with combinatorial sets of mutations that can be used to measure epistasis^11,12^. Also available are convenient approaches to measure Darwinian fitness of the mutant libraries^10,13–15^. Coupling these two technologies allows the creation of extensive genotype-to-fitness maps, or fitness landscapes^16^, that provide information important for predicting adaption^17–19^. However, these maps alone are often not sufficient to predict evolution because their topographies can depend on abiotic environmental conditions^20–23^ and biotic interactions^24,25^. Here we take two significant steps forward in fitness landscape research. First, we build on the observation that landscape structures depend on species interactions by studying the interdependence of two species’ landscapes and how they shift during coevolution. Secondly, we test whether these shifts facilitate the evolution of a key innovation.

As a model system of coevolution, we studied the host-parasite interactions among bacteriophage λ and its host, *Escherichia coli*, because of the extensive background research completed on their coevolution and the availability of well-developed molecular tools^26,27^. When λ and *E. coli* are cocultured in the laboratory, one quarter of the λ populations evolve to use a new receptor^26^. λ’s native receptor is *E. coli’s* outer-membrane protein LamB, but through mutations in its host-recognition gene *J, λ* evolves to use a second receptor protein, OmpF. While only four mutations are necessary for OmpF^+^ function^27^, more *J* mutations typically evolve along the way^26^. λ gains this new function after *E. coli* evolves resistance through *malT* mutations^26^ that cause reduced LamB expression^28^. Thus, it was hypothesized that the evolution of resistance in *E. coli* deformed λ’s fitness landscape in ways that promoted λ’s innovation^29^. In line with this, it was previously shown that some λ genotypes have higher relative fitness when cultured with resistant *malT*^−^ cells rather than ancestral cells^30^, suggesting that the host’s coevolution may promote key steps in λ’s evolution. However, not all genotypes were consistent with this pattern, and overall, too few genotypes were assayed to test the hypothesis. Here we build on this study with high throughput technologies capable of measuring the fitness of hundreds of λ genotypes in the presence of each host. This allows us to establish the contours of λ’s adaptive landscape and to determine whether host-induced deformations that naturally arise during coevolution promote OmpF^+^ evolution. Although it has been shown that antagonistic coevolution can hasten molecular evolution of phages^31^ and lead them to broader host ranges^32^, coevolution’s role in unlocking unexplored regions of the fitness landscape has not been directly tested.

## Results

### λ’s fitness landscape at different stages of coevolution

To construct λ’s fitness landscape, we focused on ten *J* mutations that were a subset of mutations λ repeatedly evolved on its path to use OmpF^26^ (Supplementary Table 1). Together they form a ten-dimensional genotype space with a total of 1,024 (2^10^) unique variants of different combinations of the mutations including the wild type allele configuration. Using Multiplexed Automated Genome Engineering (MAGE)^33^, a technique that uses repeated cycles of homologous recombination in the λ-red system to produce combinatorial genomic diversity, we were successful at engineering a library of 671 genotypes out of the possible 1,024 (see Methods). To measure the fitness of each genotype in this library, we competed the full library *en masse* and monitored their frequency changes using next generation sequencing (Supplementary Fig. 1)^34,35^. The fitness of each genotype was then computed by comparing its change in frequency relative to the non-engineered ancestor. Fitness was measured in four replicate competitions for both the ancestral host and *malT*^−^ host (see Methods). To reduce the effect of sequencing errors and to overcome other methodological pitfalls, we modified the MAGE protocol by introducing neutral *watermark* mutations in the library construction and developed a high-throughput competition assay that yielded reproduceable results (see Methods, Supplementary Fig. 2, Supplementary Discussion). Overall, we were able to measure the fitness of 580 λ genotypes cocultured with ancestral *E. coli* and 131 genotypes with *malT*^−^ (Supplementary Fig. 3). The reduction in the number of genotypes we were able to measure out of 671 in the initial library was mainly due to unfit genotypes’ frequencies falling below our limit of detection during the competitions.

Visual inspection of the two fitness landscapes reveals host-dependent structures; the landscape with the ancestral host has a standard diminishing return pattern^14,15,36,37^, while the landscape with *malT*^−^ host has an atypical sigmoidal shape that plateaus at a higher fitness than the first (Fig. 1a and 1b). The nonlinear relationship between mutation number and fitness suggests the presence of epistasis (mutation-by-mutation interactions), the differences in the magnitude of fitness effects between landscapes suggests mutation-by-host interactions, and different shapes suggest host-dependent epistasis (mutation-by-mutation-by-host interactions). To determine how much variation in fitness is explained by these interactions, we performed multiple linear regression analyses (see Methods). We found pervasive epistasis in both landscapes (Fig. 1c). For the ancestral landscape, 58.66% of the variation was explained by the direct effects of mutations and 24.69% by pairwise interactions (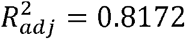, *F*_55,439_□ = 39.97, *P*□<□0.0001). Similarly, 48.35% of the variance in the *malT*^−^ landscape was explained by the direct mutation effects and 27.61% by the interaction terms (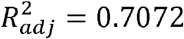, *F*_55,252_□ = 14.48, *P*□<□0.0001). To test for mutation-by-mutation-by-host interactions we regressed another linear model that includes host as a predictor variable. In this model, we found significant mutation-by-host interactions (Fig. 1d), and sizeable effects of the host-dependent epistasis (21 mutation-by-mutation-by-host interaction terms were significant out of 45 and 12.62% of the total variance in the data were attributable to these terms, Fig. 1d). This three-way interaction term measures the extent to which the landscape structure is transformed by host evolution and suggests that λ’s evolutionary trajectory could depend on its host’s genotype.

**Fig. 1.**
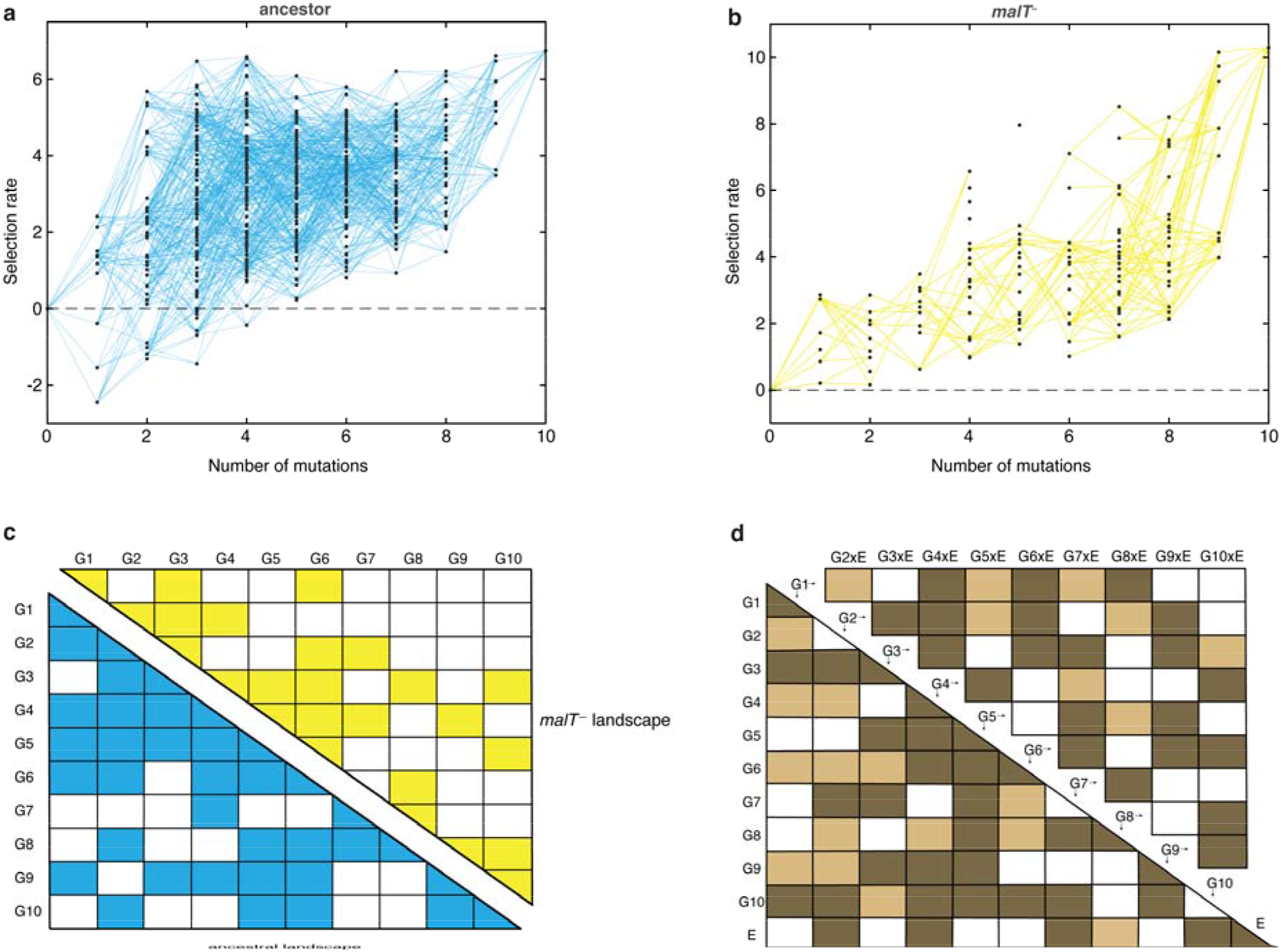
Empirical fitness landscapes of λ when infecting the a) ancestral host and b) *malT*^−^ host, and their statistical analyses in c) and d). Each node in (a) and (b) represents a unique genotype and two nodes are connected by edges if the corresponding genotypes are separated by one mutation. The node at zero mutations is ancestral λ. Selection rate (per four-hour competition experiment) is the difference of Malthusian growth rates of a given genotype *i* to ancestral λ over four hours, calculated as In (λ_*i*,4_ / λ_*i*,0_) – ln (λ_*anc*,4_ / λ_*anc*,0_) where *λ_i,t_* denotes the density of the given genotype at time *t*. **c)** Statistical analysis of direct and interactive effects of mutations in both the landscapes. Colored cells represent statistically significant terms determined by multiple regression analysis after correction for multiple hypothesis testing (see Methods). The diagonal elements of the matrix represent single mutation effects and all the off-diagonal terms represent pairwise epistatic interactions. See Supplementary Table 9 for identity of mutations corresponding to different *G_i_*. **d)** Statistical test of whether the two landscapes varied in topology. The additional variable, *E*, represents environment (host) to indicate mutation-by-host effects in the lower-left matrix and mutation-by-mutation-by-host (GxGxE) in the upper-right matrix. Light colored cells indicate terms present in the final AIC-optimized model out of the full-factorial model (*F*_76,726_ = 37.45, *P*□<□10.0001), and dark colored cells indicate statistically significant terms after controlling for rate of false positives (see Methods).

### The role of shifting landscapes in promoting λ’s innovation

To test whether changes in the structure of λ’s landscape opened trajectories to OmpF exploitation, we simulated λ’s evolution on the landscapes using a modified Wright-Fisher model (see Methods). Before running the simulations, we imputed the missing λ genotypes’ fitnesses to complete the landscapes. We did this by successively choosing missing genotypes at random and assigning them the average fitness of their nearest neighbours. The simulations were run based on conservative estimates of the number of generations and population sizes from the experiment (960 generations; ~6.3×10^9^ λ particles, Supplementary Fig. 4) and λ’s intrinsic mutation rate (7.7×10^-8^ base^-1^ replication^-1^)^38^. We predicted that λ would be more likely to evolve OmpF function in simulations that accounted for coevolution by shifting the population from one landscape to the next. We ran trials where λ evolved on only one landscape at a time to establish a baseline for the frequency of OmpF^+^ evolution without coevolution. Next, we ran nine shifting landscape scenarios where we varied how many generations λ evolved on the ancestral host landscape before switching to the *malT*^−^ landscape. As anticipated, the switching protocol increased the frequency of OmpF^+^ evolution in all 9 treatments above the single host simulations, but only 7 out of 9 treatments were found to be significantly higher (Fig. 2; ANOVA: F-ratio = 6.14, *d.f*. = 99, *P* < 0.0001, Supplementary Table 2). This result was robust to changes in population size and total number of generations, and when controlling for both, different number of genotypes measured in the two landscapes, and noise created by imputing missing data points (Supplementary Fig. 5, Supplementary Discussion).

**Fig. 2.**
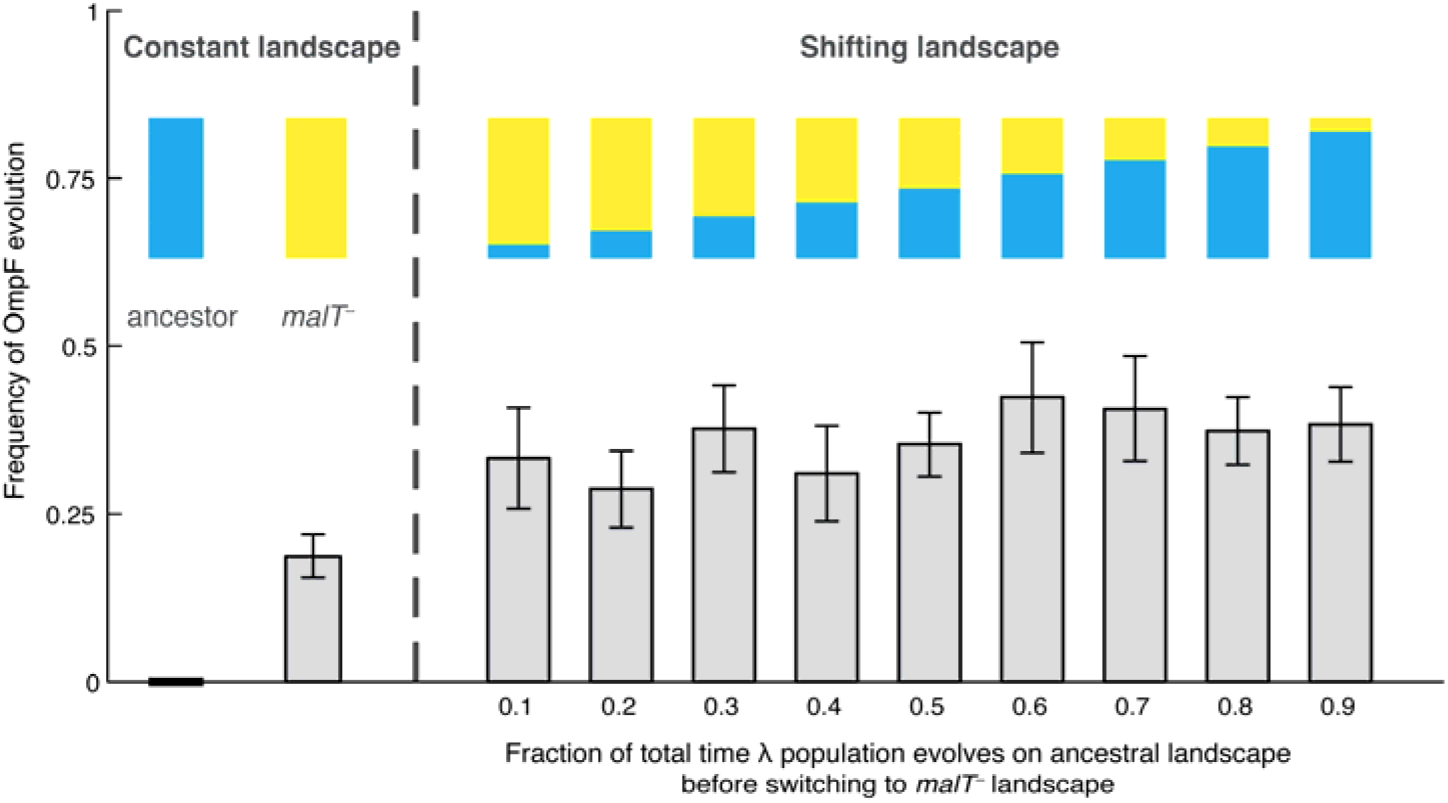
Frequency of OmpF innovation observed when λ evolution is simulated on shifting fitness landscapes. Each bar represents an average of 300 simulation runs. Error bars indicate 95% confidence intervals. OmpF evolution is favoured when λ evolves on shifting landscapes. The only two shifting landscape treatments that are not significantly higher than simulations on the constant *malT*^−^ landscape are the 0.2 and 0.4 treatments (Supplementary Table 2).

### Reconstructing coevolution in an experimental population

The simulation results suggest that the shifting landscape encourages λ’s evolution to OmpF function. In particular, the simulations show that the first steps along the path to the innovation are more likely if λ first adapts to the ancestral bacterium, meanwhile the final steps are more likely to occur if the host coevolves resistance. To verify this result with laboratory experiments, we analyzed the path λ took to OmpF^+^ in a single population cryopreserved from the previous coevolution study (population ‘D7’ in Meyer *et al*. 2012^(26)^; Table S1). We sampled λ strains from different timepoints and sequenced their *J* gene (Fig. 3a, Supplementary Table 3 and Supplementary Table 4). Next, we ran pairwise competition experiments between λ genotypes at different stages of evolution on the two hosts. We found that the first mutation on the line of descent to OmpF^+^ required ancestral *E. coli* to evolve, while the second mutation required *malT*^−^ *E. coli* (Fig. 3b and 3c). In addition, the OmpF^+^ genotype with five *J* mutations only outcompeted the genotype with two mutations when provided with *malT*^−^ hosts (Supplementary Fig. 6). These findings show that the path λ took in population D7 required it to sequentially adapt to both host types and that λ’s fitness landscape changed during coevolution in a way that ultimately facilitated evolutionary innovation.

**Fig. 3.**
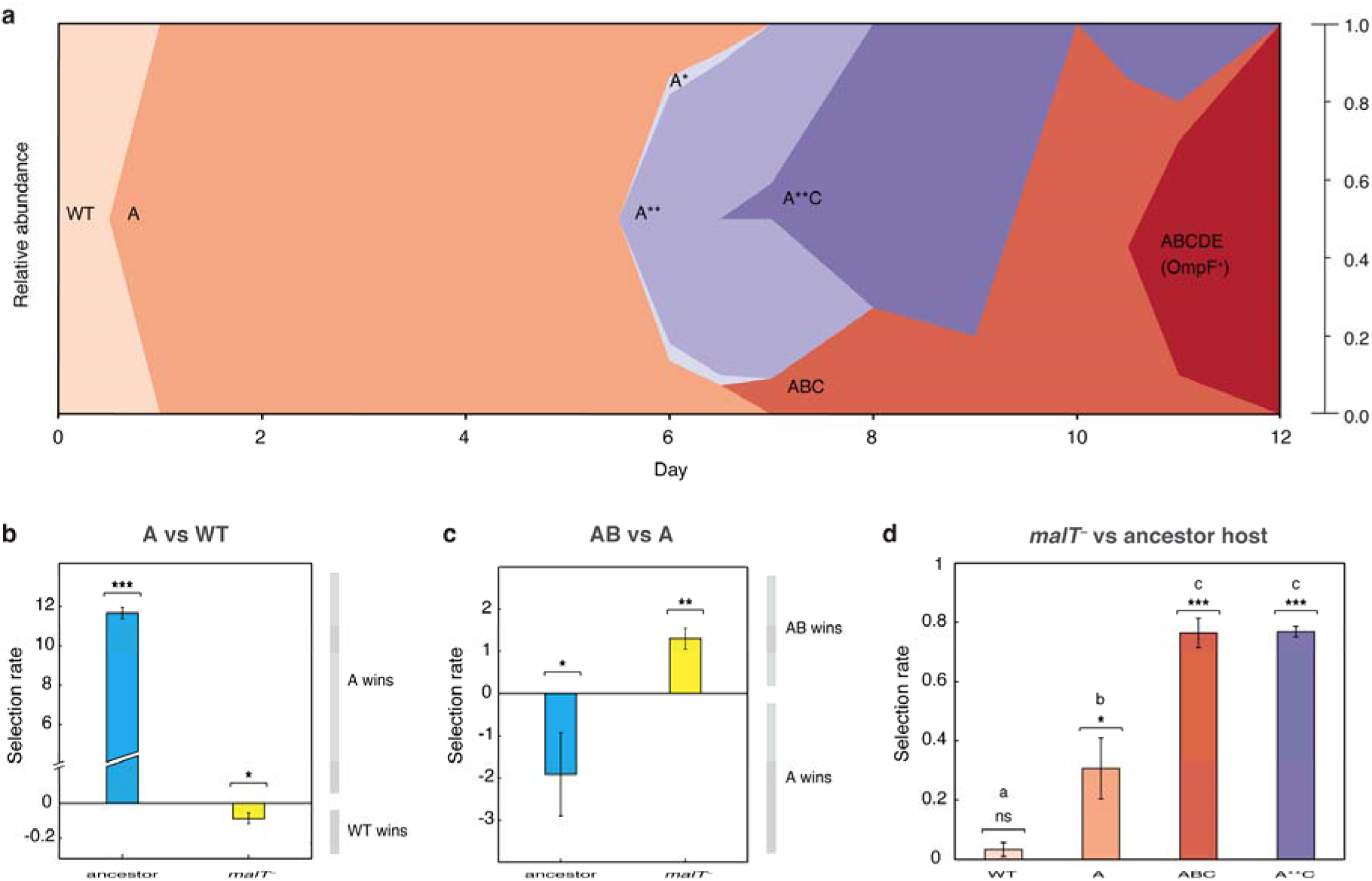
*J* evolution (a) and evidence of the interdependency between λ and *E. coli* fitness during their coevolution (b-d). **a)** Phylogenetic reconstruction and relative abundance of λ genotypes isolated through time from a previously coevolved community^26^. Each letter and star indicate a non-synonymous mutation in *J* (see Supplementary Table 3 for labels’ corresponding mutations). A genotype’s relative abundance on a given day is denoted by the fraction of the total height of the y-axis that it occupies (e.g., on Day 9, frequency of ABC is 0.2 and A**C is 0.8; see Supplementary Table 4). The lineage WT-A-ABC-ABCDE eventually evolves OmpF function and fixes in the population; resistance in *E. coli* through *malT* rises to high frequencies between days five and eight^26^. **b) & c)** Selection rates (per 24 h) of phage genotypes on the two hosts. Each bar represents the mean of three experimental replicates. While mutation (A) is favored over wildtype λ in the presence of the ancestral host and not *malT*^−^ AB only outcompetes A in the presence of *malT*^−^ and not the ancestral host. One tailed t-tests to test if the mean selection rate is significantly greater (or less) than zero: A vs WT with ancestor host- *t* = 98.76, *d.f*. = 2, *P* < 0.0001; A vs WT with *malT*^−^ - *t* = −4.99, *d.f*. = 2, *P* = 0.0190; AB vs A with ancestor- *t* = 3.4, *d.f*. = 2, *P* = 0.0383; AB vs A with *malT*^−^ - *t* = −8.88, *d.f*. = 2, *P* = 0.0062. **d)** Selection rate (per 4 h) of *malT*^−^ *E. coli* relative to its ancestor in the presence of λ from different stages of coevolution. Each competition was replicated three times. Lowercase letters denote significance via Tukey’s honest significance test, see Supplementary Table 7 for pairwise P-values (ANOVA: F-ratio = 111.22, *d.f*. = 11, *P* < 0.0001). One tailed t-tests were also used to test if the selection rate of *malT*^−^ was greater than zero-WT: *t* = 2.44, *d.f*. = 2, *P* = 0.676: A: *t* = 5.12, *d.f*. = 2, *P* = 0.0181; ABC: *t* = 26.59, *d.f*. =2, *P* = 0.0007; A**C: *t* = 71.67, *d. f*. = 2, *P* <0.0001. This shows that *malT*^−^ is unlikely to evolve in the presence of WT λ but it becomes progressively more likely as λ gains mutations. Asterisks over all the competitions indicate significance level corresponding to the P-values. Error bars in all bar graphs represent one sample standard deviation.

At this point, we shifted the focus of our studies to testing whether *E. coli’s* resistance evolution was also impacted by λ’s evolution. Before reconstructing *J* evolution, it was believed that *E. coli* evolved resistance and then *J* mutations evolved in response^26^. However, *J* evolved within a day, while it was previously shown that *malT*^−^ mutations take about a week to evolve^26^. The timing suggests that λ improved infectivity and then applied pressure on *E. coli* to evolve resistance. To test whether λ evolution promoted host resistance evolution, we ran competition experiments between ancestral and *malT*^−^ hosts in the presence of phages isolated from four different time points. We found that *malT*^−^ was not significantly more fit than the ancestral *E. coli* in the presence of the ancestral λ, but it was more fit in the presence of the evolved λs (Fig. 3d). This result combined with the others suggests that there is an intricate coevolutionary feedback at play between λ and *E. coli*: λ evolves *J* mutations that better exploit *E. coli*, which in turn applies pressure on *E. coli* to evolve resistance. Once resistance evolves, new adaptive pathways become available to λ that encourage the innovation. For the computer simulations, we arbitrarily chose timepoints to switch from one host to the other; however, in reality, the dynamics of the switch are dictated by the host-parasite coevolution.

### Evolutionary replay experiments

To further test the role of coevolutionary processes at driving λ’s innovation, we ran replays of the coevolution experiment (Supplementary Fig. 7). We initiated 12 populations with *malT*^−^ host that already possessed resistance, and 12 populations with ancestral host where λ and *E. coli* would coevolve normally. The former treatment should hinder the evolution of OmpF function because it denies λ the opportunity to evolve first with ancestral *E. coli*. In line with our expectations, 0 of 12 replicates evolved OmpF use in the *malT*^−^ treatment, meanwhile 3 out of 12 evolved the innovation in the ancestral treatment (Fig. 4a and 4b). By sequencing *J* alleles of the resulting λ genotypes, we found that fewer mutations evolved with *malT*^−^ despite evolving for the same length of time. This suggests that λ’s evolution was stymied by starting with the resistant host (Supplementary Table 5), and by disrupting the coevolutionary process we interfered with λ’s ability to innovate.

**Fig. 4.**
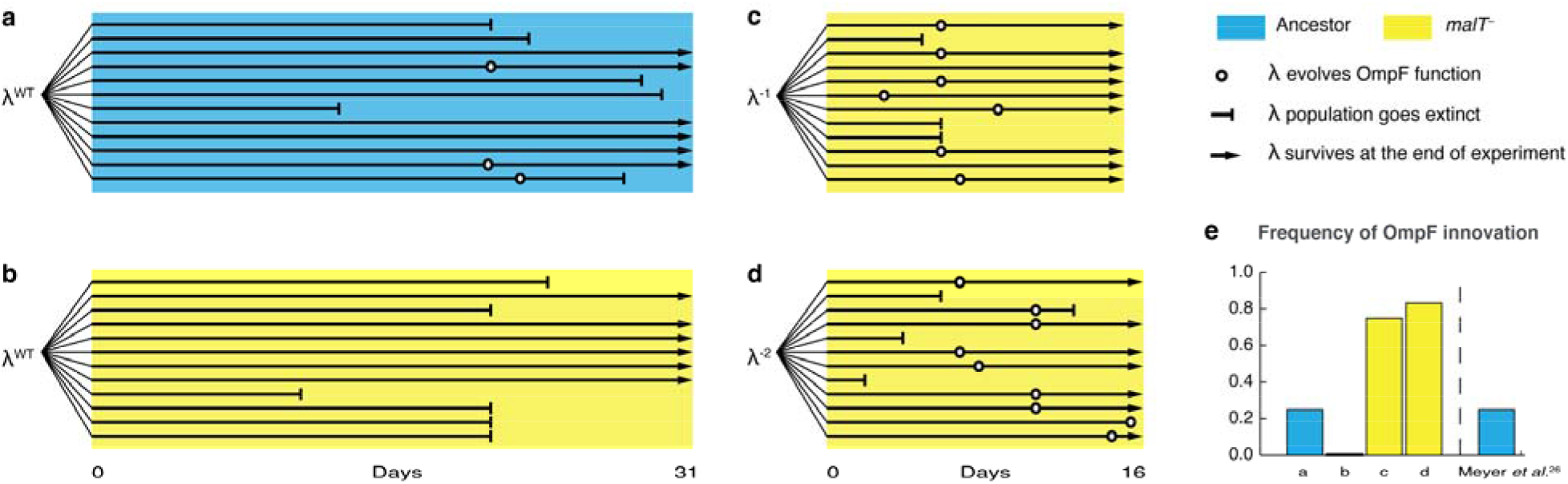
Evolutionary replay experiments reveal that λ’s evolution to use OmpF depends on host coevolution. **a)** Wildtype λ with ancestral host, **b)** wildtype λ with *malT*^−^, **c)** λ one mutation removed from evolving OmpF function with *malT*^−^, and **d)** identical setup as (c), but with λ two mutations removed (see Supplementary Table 6 for identity of mutations). **e)** The bar graph provides the frequency of OmpF^+^ evolution compared to the frequency observed by Meyer *et al*.^26^. Given λ’s established 1 in 4 rate of OmpF^+^ with ancestral host, the probability of observing no OmpF^+^ evolution in 12 replicate populations is ~0.03. Thus, no positives for OmpF evolution in (b) shows that λ’s evolution to OmpF function is significantly hindered when coevolution is initiated with *malT*^−^ host. However, *malT*^−^ does not impede OmpF^+^ evolution when the coevolution is initiated with already evolved λs (P-values for Fisher’s exact test: between (b) and (c)- *P* = 0.0261; between (b) and (d)- *P* = 0.0122). Notably, some λ populations went extinct which is common for these experiments and was previously shown to be caused by the evolution of resistance mutations in *E. coli*’s ManXYZ protein complex^26^.

Lastly, we tested whether *malT*^−^ would still promote the evolution of OmpF use if the replay experiments were initiated with λ genotypes positioned further along the path to OmpF^+^. We initiated two more replays: one with a λ strain that was just a single mutation away from OmpF^+^, and another that was two mutations away (see Supplementary Table 6 for J alleles). 16 of 24 λ populations evolved OmpF^+^ showing that whether *malT*^−^ promotes OmpF^+^ evolution depends on where the λ genotypes are located in the fitness landscape (Fig. 4c and 4d).

## Discussion

This study provides a fuller and more complex understanding of λ and *E. coli* coevolution than previously published. λ was thought to evolve OmpF use as a direct response to *malT*^−^ resistance^26^; however, here we learned that key steps were missing from that model. λ is initially poor at infecting its host, so it evolves mutations in *J* that enhance its infectivity. The new λ genotypes apply pressure on *E. coli* to evolve resistance. When host-resistance increases in the community, λ’s fitness landscape is deformed in a way that promotes J evolution towards OmpF use, but only if it had already acquired some J mutations. Remarkably, the timing and coordination of each of these interdependent steps is facilitated by the reciprocity ingrained in host-parasite coevolution.

Taken together, our studies show that the fitness of a parasite depends on complex genetic interactions within its own genome and with the genomes of interacting hosts. These interdependencies result in highly contingent evolution where λ is unlikely to evolve an innovation unless it participates in a particular sequence of coevolutionary steps with its host. Despite the stochasticity that is expected to arise in systems with substantial historical contingency^39,40^, λ’s evolution to use a new receptor is repeatable because the sequence is coordinated by coevolutionary feedbacks. While coevolution may yield tangled banks of interactions, we demonstrate how high throughput technologies can be used to untangle them and to predict evolution despite their complexity. With this approach, we were able to provide direct experimental evidence that fluctuating landscapes, also known as fitness seascapes^41^, can promote evolutionary innovations.

This work was completed during the 2020 SARS-CoV-2 pandemic, raising the question of whether this research provides insight into strategies to prevent future pandemics. One existing recommendation to prevent emerging infectious diseases is to limit human incursion into natural places in order to reduce exposure to zoonotic diseases^42,43^. Our research suggests that human incursions and other environmental perturbations may also enhance the chance that exposure will result in a host shift. Subjecting viruses to changing environmental conditions may favour the evolution of new strains with increased potential to shift.

## Acknowledgments

We thank the James McDonnel Foundation and the NSF-DEB (1934515) for financial support and The Max Planck Society for supporting JG. We thank A. Agarwal for help with the statistical analyses of fitness landscapes.

## Author Contributions

AG conducted laboratory experiments, ran computational simulations, created all the figures, and wrote the paper, LZ helped develop the project and with computational analyses, HMS, JG, and ARB conducted experiments, BK oversaw a portion of the studies, ET helped with simulations, RK helped with technology development and oversaw aspects of the work, JRM collected MAGE-Seq data, wrote the manuscript, and oversaw the project, and all coauthors edited the text.

## Competing interests

The authors declare no competing interests.

**Correspondence and requests for materials** should be addressed to R.K. or J.R.M.

## Data and code availability

Amplicon sequencing data used to generate fitness landscapes in this study have been deposited in the NCBI Sequence Read Archive under BioProject PRJNA646809. Data that support the findings of this study will be deposited at Dryad, and code will be made available at GitHub on publication.

## Notes

### Competing Interest Statement

The authors have declared no competing interest.

## References

1 Orr, H. A. The genetic theory of adaptation: a brief history. Nat Rev Genet 6, 119–127, doi:10.1038/nrg1523 (2005).

2 Gavrilets, S. in Evolution, the extended synthesis (eds Massimo Pigliucci & Gerd Müller) Ch. 3, 45–79 (MIT Press, 2010).

3 Van Valen, L. A new evolutionary law. Evolutionary Theory 1, 1–30 (1973).

4 Doebeli, M. Adaptive diversification. (Princeton University Press, 2011).

5 Thompson, J. N. The geographic mosaic of coevolution. (University of Chicago Press, 2005).

6 Nahum, J. R. et al. Improved adaptation in exogenously and endogenously changing environments. The 2019 Conference on Artificial Life, 306–313, doi:10.1162/isal_a_052 (2017).

7 Thompson, J. N. & Cunningham, B. M. Geographic structure and dynamics of coevolutionary selection. Nature 417, 735–738, doi:10.1038/nature00810 (2002).

8 Zaman, L. et al. Coevolution Drives the Emergence of Complex Traits and Promotes Evolvability. PLOS Biology 12, e1002023, doi:10.1371/journal.pbio.1002023 (2014).

9 Darwin, C. The Origin of Species. (John Murray, 1859).

10 Weinreich, D. M., Delaney, N. F., Depristo, M. A. & Hartl, D. L. Darwinian evolution can follow only very few mutational paths to fitter proteins. Science 312, 111–114, doi:10.1126/science.1123539 (2006).

11 Kosuri, S. & Church, G. M. Large-scale de novo DNA synthesis: technologies and applications. Nature Methods 11, 499–507, doi:10.1038/nmeth.2918 (2014).

12 Fowler, D. M. & Fields, S. Deep mutational scanning: a new style of protein science. Nat Methods 11, 801–807, doi:10.1038/nmeth.3027 (2014).

13 Palmer, A. C. et al. Delayed commitment to evolutionary fate in antibiotic resistance fitness landscapes. Nature Communications 6, 7385, doi:10.1038/ncomms8385 (2015).

14 Khan, A. I., Dinh, D. M., Schneider, D., Lenski, R. E. & Cooper, T. F. Negative epistasis between beneficial mutations in an evolving bacterial population. Science 332, 1193–1196, doi:10.1126/science.1203801 (2011).

15 Chou, H. H., Chiu, H. C., Delaney, N. F., Segre, D. & Marx, C. J. Diminishing returns epistasis among beneficial mutations decelerates adaptation. Science 332, 1190–1192, doi:10.1126/science.1203799 (2011).

16 Wright, S. The roles of mutation, inbreeding, crossbreeding, and selection in evolution.Vol. 1 (na, 1932).

17 de Visser, J. A. & Krug, J. Empirical fitness landscapes and the predictability of evolution. Nat Rev Genet 15, 480–490, doi:10.1038/nrg3744 (2014).

18 de Visser, J. A. G. M., Elena, S. F., Fragata, I. & Matuszewski, S. The utility of fitness landscapes and big data for predicting evolution. Heredity (Edinb) 121, 401–405, doi:10.1038/s41437-018-0128-4 (2018).

19 Lee, J. M. et al. Deep mutational scanning of hemagglutinin helps predict evolutionary fates of human H3N2 influenza variants. Proc Natl Acad Sci U S A 115, E8276–E8285, doi:10.1073/pnas.1806133115 (2018).

20 Ogbunugafor, C. B., Wylie, C. S., Diakite, I., Weinreich, D. M. & Hartl, D. L. Adaptive Landscape by Environment Interactions Dictate Evolutionary Dynamics in Models of Drug Resistance. PLoS Comput Biol 12, e1004710, doi: 10.1371/journal.pcbi.1004710 (2016).

21 Lindsey, H. A., Gallie, J., Taylor, S. & Kerr, B. Evolutionary rescue from extinction is contingent on a lower rate of environmental change. Nature 494, 463–467, doi:10.1038/nature11879 (2013).

22 Steinberg, B. & Ostermeier, M. Environmental changes bridge evolutionary valleys. Sci Adv 2, e1500921, doi:10.1126/sciadv.1500921 (2016).

23 Flynn, K. M., Cooper, T. F., Moore, F. B. & Cooper, V. S. The environment affects epistatic interactions to alter the topology of an empirical fitness landscape. PLoS Genet 9, e1003426, doi:10.1371/journal.pgen.1003426 (2013).

24 Cervera, H., Lalić, J. & Elena, S. F. Effect of Host Species on Topography of the Fitness Landscape for a Plant RNA Virus. J Virol 90, 10160–10169, doi:10.1128/JVI.01243-16 (2016).

25 Fragata, I., Blanckaert, A., Dias Louro, M. A., Liberles, D. A. & Bank, C. Evolution in the light of fitness landscape theory. Trends Ecol Evol 34, 69–82, doi:10.1016/j.tree.2018.10.009 (2019).

26 Meyer, J. R. et al. Repeatability and contingency in the evolution of a key innovation in phage lambda. Science 335, 428–432, doi:10.1126/science.1214449 (2012).

27 Maddamsetti, R. et al. Gain-of-function experiments with bacteriophage lambda uncover residues under diversifying selection in nature. Evolution 72, 2234–2243, doi:10.1111/evo.13586 (2018).

28 Boos, W. & Bohm, A. Learning new tricks from an old dog: MalT of the Escherichia coli maltose system is part of a complex regulatory network. Trends Genet 16, 404–409, doi:10.1016/S0168-9525(00)02086-2 (2000).

29 Thompson, J. N. Evolution. The role of coevolution. Science 335, 410–411, doi:10.1126/science.1217807 (2012).

30 Burmeister, A. R., Lenski, R. E. & Meyer, J. R. Host coevolution alters the adaptive landscape of a virus. Proc Biol Sci 283, doi:10.1098/rspb.2016.1528 (2016).

31 Paterson, S. et al. Antagonistic coevolution accelerates molecular evolution. Nature 464, 275–278, doi:10.1038/nature08798 (2010).

32 Hall, Alex R., Scanlan, Pauline D. & Buckling, A. Bacteria□Phage Coevolution and the Emergence of Generalist Pathogens. The American Naturalist 177, 44–53, doi:10.1086/657441 (2011).

33 Wang, H. H. et al. Programming cells by multiplex genome engineering and accelerated evolution. Nature 460, 894–898, doi:10.1038/nature08187 (2009).

34 Kelsic, E. D. et al. RNA Structural Determinants of Optimal Codons Revealed by MAGE-Seq. Cell Syst 3, 563–571 e566, doi:10.1016/j.cels.2016.11.004 (2016).

35 Russ, D. et al. Escape mutations circumvent a tradeoff between resistance to a betalactam and resistance to a beta-lactamase inhibitor. Nat Commun 11, 2029, doi:10.1038/s41467-020-15666-2 (2020).

36 Kryazhimskiy, S., Rice, D. P., Jerison, E. R. & Desai, M. M. Microbial evolution. Global epistasis makes adaptation predictable despite sequence-level stochasticity. Science 344, 1519–1522, doi:10.1126/science.1250939 (2014).

37 MacLean, R. C., Perron, G. G. & Gardner, A. Diminishing Returns From Beneficial Mutations and Pervasive Epistasis Shape the Fitness Landscape for Rifampicin Resistance in *Pseudomonas aeruginosa*. Genetics 186, 1345, doi:10.1534/genetics.110.123083 (2010).

38 Drake, J. W. A constant rate of spontaneous mutation in DNA-based microbes. Proc Natl AcadSci U S A 88, 7160–7164, doi:10.1073/pnas.88.16.7160 (1991).

39 Gould, S. J. Wonderful life: the Burgess Shale and the nature of history.1st edn, (W.W. Norton, 1989).

40 Blount, Z. D., Lenski, R. E. & Losos, J. B. Contingency and determinism in evolution: Replaying life’s tape. Science 362, doi:10.1126/science.aam5979 (2018).

41 Merrell, D. J. The adaptive seascape: the mechanism of evolution. (University of Minnesota Press, 1994).

42 Jones, K. E. et al. Global trends in emerging infectious diseases. Nature 451, 990–993, doi:10.1038/nature06536 (2008).

43 Allen, T. et al. Global hotspots and correlates of emerging zoonotic diseases. Nature Communications 8, 1124, doi:10.1038/s41467-017-00923-8 (2017).

