## Supplementary material for "Host-parasite coevolution promotes innovation through deformations in fitness landscapes": Methods and Supplementary Information

### Methods and Supplementary Information for

**This file includes:**

Methods
Supplementary Discussion
Supplementary Figures 1 to 8
Supplementary Tables 1 to 12

#### Methods

Bacterial and phage strains

The ancestral phage λ strain in our study is cI26; it is a strictly lytic strain that was used in the previous coevolution study on which this paper builds[1]. λ has two life cycles; lytic and lysogenic. In the lytic life cycle, λ infects its host, creates multiple copies of its genome, and lyses the cell to produce new viral particles. This is unlike the lysogenic life cycle of λ, where it stably integrates its genome into the host genome and is replicated along with the host[2]. The λ strain cI26 has a frameshift mutation in the *cI* gene that disrupts the regulatory protein cI required by λ to switch to a lysogenic life cycle[1]. This renders cI26 obligatory lytic. The other λ strain we used was cI857 (provided by Ing-Nang Wang, State University of New York at Albany) to construct our λ genomic library using MAGE (discussed in 'Fitness landscapes' section). The advantage to using cI857 over cI26 is that it is able to form a lysogen and as a genomic complex with *E. coli*'s genome, its genome can be easily edited with *E. coli* engineering methods. cI857 has an advantage over typical lysogenic λ because it has a mutation in the *cI* gene that makes the repressor protein cI unfold at high temperatures[3]. This enables us to induce lytic life cycle of cI857 from lysogens using heat shock treatments, which has fewer side effects compared to using mutagens that the typical strain requires for induction.

We used *Escherichia coli* B strain REL606 for the ancestral-sensitive host and its derivative EcC4 for the evolved-resistant (*malT*⁻) host to construct fitness landscapes (in Fig. 1) and run competition assays (in Fig. 3). REL606 was the ancestral host in the coevolution experiments performed by Meyer *et al.*[1], and EcC4 was isolated from one of the coevolving populations that has a single mutation, a nonsense mutation (C→T) at genome location 3,482,567 in *malT* gene (Table S5 in Meyer *et al.*[1]). MalT is a positive regulator of *lamB*, so a disruption in transcription of *malT* inhibits LamB expression[4]. Resistance to λ in EcC4 due to this nonsense *malT* mutation has been shown to yield high levels of resistance, the equivalent of a nonsense mutation in *lamB* (Fig. S5 in Chaudhry *et al.*[5]). We used another *malT*⁻ mutant of REL606, named LR01, for our coevolutionary replay experiments (Fig. 4). This strain has a 25-bp duplication at genome location 3,482,677 causing a frameshift in *malT*. LR01 was also isolated from a coevolving population of λ and *E. coli* (Fig. S3 in Meyer *et al.*[1]), however, unlike EcC4, LR01 has a high reversion rate to *malT*⁺ that results in "leaky resistance" to λ[(5)]. This allows λ populations to sustain serial dilution transfers, and thus was critical for the success of the coevolutionary replay experiments (Fig. 4).

A number of *E. coli* strains were used for culturing λ. The most often used was DH5α, a $lacZ\alpha^-$ derivative of *E. coli* K-12, because it is permissive to all λ genotypes and it lacks *lacZ* which is used as a genetic marker to distinguish phage genotypes in competition assays (additional information in 'Phage Competition Experiments' section). Two *lamB*⁻ mutants of *E. coli* with nonfunctional LamB were used interchangeably to culture OmpF⁺ λ on Petri dishes. One strain was a derivative of REL606 that has a 1-bp insertion of nucleotide T between base positions 610 and 611 in *lamB*[1,6], and the other strain was an *E. coli* K-12 derivative from the Keio collection (*lamB*⁻ JW3996)[7]. We found no difference between the efficiency of plaquing on two strains.

Multiplexed Automated Genome Engineering (MAGE) was performed in an *E. coli* K12 strain, HWEC106, provided by Harris Wang, Columbia University. The strain's *mutS* gene is deleted and it possesses the pKD46 plasmid with an inducible λ-red recombineering system[8].

cI857 successfully integrated into this strain's canonical ATTB site located genomically near the *galK* gene[1,9,10].

Media

We performed most experiments in conditions identical to the initial coevolution experiment[1]. For competition assays (low and high throughput) and two of the four coevolutionary replay experiments (Fig. 4a and 4b), bacteria and phage were cocultured in **modified M9 Glucose** (47.7 mM disodium phosphate, 22.0 mM potassium phosphate monobasic, 18.7 mM ammonium chloride, 8.6 mM sodium chloride, 0.1 mM calcium chloride, 10 mM magnesium sulfate and 5.55 mM glucose) as used in Meyer *et al.*[1]. A new medium was used for the remaining two coevolution replays (Fig. 4c and 4d) that we call **Tris DM Glucose**: 1.6 mM potassium phosphate monobasic, 0.59 mM potassium phosphate dibasic, 0.2 mM of calcium chloride, 50 mM Tris base (pH 7.4), 10 mM of magnesium sulphate, 7.5 μM thiamine, 3.2 mM ammonium sulphate, and 5.55 mM glucose. This medium is improved over M9 Glucose because it is less prone to magnesium precipitation, and fortunately has no noticeable effect on the coevolution.

For isolation, estimating densities and initial culturing of bacteria and phage, four more media were used according to the following specifications— a) **LB** (Lennox Broth): 10 g tryptone, 5 g yeast extract, and 5 g sodium chloride per liter of water, b) **LBM9**: 20 g tryptone, 10 g yeast extract, 12.8 g sodium phosphate heptahydrate, 3 g potassium phosphate monobasic, 0.5 g sodium chloride, 1 g ammonium chloride, 1.2 g magnesium sulfate, 22 mg calcium chloride per liter of water, c) **LB agar**: 10 g tryptone, 5 g yeast extract, 5 g sodium chloride, and 16 g agar per liter of water, d) **Soft agar**: 10 g tryptone, 1 g yeast extract, 8 g sodium chloride, 7 g agar, 0.1 g glucose per liter of water, supplemented with a final concentration of 2 mM calcium chloride. The soft agar was also supplemented with 10 mM magnesium sulphate to improve plaquing of phage particles.

Isolation and culturing techniques

Most strains were grown at 37°C, except for HWEC106, which was grown at 30°C because pKD46 has a temperature sensitive origin of replication. To ensure uniform aeration and nutrient availability, 4 ml cultures were shaken at 220 rpm and 10 ml cultures at 120 rpm; different rpm was chosen to ensure equivalent aeration in cultures (10 ml cultures were grown in 50-ml Erlenmeyer flasks and 4ml in glass tubes). We used LBM9 to grow phage cultures, and LB to grow bacteria cultures, unless otherwise indicated. All bacteria and phage stocks were stored by freezing 1 ml of culture with 15% glycerol at -80°C. We revived phage from freezer stocks by growing ~2 μl of frozen stocks on 100 μl of DH5α overnight culture in 4 ml of LBM9. To harvest phage from this, cells were killed and separated by adding 100 μl of chloroform and centrifuging the solution at 3900 rpm for 10 min. The supernatant containing phage lysate was then stored with ~2% chloroform at 4°C. Bacteria were revived from -80°C by growing ~2 μl of the frozen stocks over night in 4 ml of LB.

Phage strains were isolated from a population by infusing phage particles into bacterial lawns of DH5α cells. We made these *infused plates* by mixing a small volume (between 10 and 100 μl) of diluted phage and ~5x10$^8$ host cells to a 4 ml of molten soft agar at 55°C, and pouring the mixture over an LB agar plate[11]. The soft agar was allowed to solidify and then incubated overnight at 37°C for phage particles to form plaques. A plaque is a near-circular clearing that forms on the bacterial lawn when nearby cells are killed by an infection that is initiated by a single phage particle. Phage dilutions were made in saline solution (8.5 g/L NaCl) and implemented to yield between 30 and 300 plaques. A single plaque was picked from the infused plates and clonal phage stocks were made from it by culturing it overnight with DH5α and extracting phage lysate from it using techniques similar to frozen λ stocks.

Estimation of phage densities

Infused plates with appropriate dilutions of phage in saline solution (8.5 g/L NaCl) were used to estimate phage densities for phage competition assays. We controlled the number of plaques on an infused plate so that individual plaques could be identified and counted. These counts were used to back-calculate phage growth rate.

To estimate the phage densities in the coevolutionary replay experiments, serial dilutions of phage were spotted on a bacterial lawn of DH5α. We made the bacterial lawns by adding ~$5x10^9$ cells of DH5α to 10 ml of molten soft agar at 55°C, and pouring it over a 150-mm diameter Petri dish of LB agar base[11]. After the soft agar solidified, 2 μl of eight different phage dilutions was added onto the surface, let dry, and incubated overnight at 37°C. The plaques were counted from the dilution where we could identify individual plaques. We then used these counts to calculate phage density in pfu per ml, where pfu is plaque forming unit.

Sequencing for analyses of mutations in J

We routinely sequenced the *J* gene of λ to verify the identity of stocks and to determine whether *J* evolved in the re-play experiments. DNA samples were PCR amplified using Q5® High-Fidelity 2X master mix (New England Biolabs) and primers described in Supplementary Table 8. Unpurified PCRs were submitted to Genewiz (La Jolla, CA) for Sanger sequencing.

Calculation of selection rate using Malthusian parameter

Selection rate ($s$) was used to quantify the difference in fitness between pairs of strains (say **X** and **Y**) in a given environment. It is calculated as the difference of their Malthusian parameters:

$$s = \frac{ln\frac{X_T}{X_0} - ln\frac{Y_T}{Y_0}}{T} \qquad (1)$$

where $X_t$ and $Y_t$ are the densities of strains **X** and **Y**, respectively, at time $t$, and $T$ is the period of time over which strains are assessed (and the assay starts at $t = 0$). Selection rate has units of inverse time, however, in all our figures, we report selection rate as per unit time period ($T$) of the assays. We reported the values this way because λ does not strictly grow exponentially and it can be misleading to report normalized rates after dividing by $T$ because the rates cannot be extrapolated for different times in a straightforward way like for exponential growths. That being said, for readers who want the per hour rates, we have provided the value of $T$ in hours within each figure legend.

Construction of λ genomic library using Multiplexed Automated Genome Engineering (MAGE)

Our goal was to measure the fitness of a significant number of λ genotypes to establish the basic structure of the landscape. To accomplish this, we generated a combinatorial library of genotypes made with 10 *J* mutations previously observed to evolve *en route* to $OmpF^+$

(Supplementary Table 1). These mutations were chosen because they fell within two <100 base frames where adaptive mutations tended to evolve. We used a genetic engineering technique called Multiplexed Automated Genome Engineering (MAGE) to construct the library[12]. This technique employs the λ-red recombineering system that can efficiently recombine single stranded DNA into *E. coli*'s genomic DNA. To engineer λ, λ's genome is integrated into *E. coli*'s genome creating a lysogen. The λ genome in the lysogen (a prophage) becomes dormant and can be treated as any other *E. coli* gene. Gene edits in this λ lysogen are made by expressing λ-red from a plasmid (pKD46) within *E. coli* cells[8] and electroporating synthetic oligos with specific mutations written into the sequences. Multiple oligos can be combined into a single experiment in order to create a diverse library of different combinations of the mutations. The output of this procedure is a single sample with a library of genotypes mixed together.

We previously reported our MAGE protocol in Maddamsetti *et al.*[9]. The oligos used are in Table S1 of the manuscript listed under the subheading '10-mutation library'. There are a few important aspects of the oligo and protocol design that we discuss here, but for more detailed information see the original paper.

Our goal was to construct a genetic library with each combination of mutations represented equally, this means a 50/50 split of mutant and wild type states at each site. To achieve this, we modified the typical MAGE protocol. First, when mutations were clustered near each other such that a single 90-mer used for MAGE editing encompassed multiple mutations, we designed multiple oligos with different combinations of the mutations at these nearby sites. Without doing this, edits would be correlated, creating an imbalance in the network representation, and limiting our ability to determine the effects of individual mutations. Secondly, we designed oligos with the wild type state at the 10 positions. Including these oligos slowed the efficiency of MAGE to introduce mutations because sometimes they would overwrite an edit, however, it safeguarded the procedure from saturating the library with the 10 mutations. A third strategy was employed to even out the variation at these ten sites. This was to perform MAGE starting from two orthogonal points in the genetic network; λ that had none of the 10 mutations and one that we engineered in all 10[(9)]. Additionally, we ran MAGE on three separate replicates from each starting point in order to enhance the potential to explore more genetic space. Lastly, we ran the MAGE protocol for 50 cycles, which is the number of cycles a computer simulation of MAGE with an efficiency of 10% recombination efficiency per cycle predicted we would need to construct all variants. In the end, we mixed all six MAGE libraries together in equal frequencies to maximize diversity in the library.

Our design also included measures to improve our ability to detect rare variants in the library. The next step of the fitness landscape protocol is amplicon sequencing with an Illumina MiSeq. This technology has relatively low rates of error (~$10^{-3}$ per base sequenced), however, we anticipated that some of the genotypes in our library would fall below this frequency, making it impossible to distinguish between false positives and true reads. To improve our resolution, we edited a synonymous mutation adjacent to each focal mutation. We called this a *watermark* mutation because it helped us distinguish between a true genome edit and error in sequencing. By only recording the presence of a focal mutation if it occurred alongside the watermark, we decreased our detection limit from ~$10^{-3}$ to ~$10^{-6}$.

The watermark strategy we employed was slightly more complicated than this. Most synonymous mutations at the C-terminal end of proteins, like our watermark mutations, have no fitness effects[13]. Despite this, we designed a way to test for neutrality. For each focal mutation we edited one of two different watermark mutations (see Supplementary Table 9 for list of mutations). This allowed us to compare equivalent genotypes with distinct watermarks in order test whether one of the synonymous mutations impacted λ's fitness when compared to the other. We found that there was no significant fitness effect of the neutral mutations on the phage genotype (see Supplementary Discussion).

Empirical fitness landscape resolved through one-pot competition experiment

After creating the λ genomic library, we measured the relative fitness of genotypes with respect to the wild type λ. To do this, we cocultured λ genomic library and ancestral λ in a 1:9 ratio. This ratio was used so that the most abundant competitor remains the ancestor throughout the competition experiment. This means that the competitive fitness we measured is with respect to a single genotype. If we had not done this, then as the community of engineered λ shifts during the competition, and more fit genotypes become enriched, they will change the mean fitness of the population and cause mildly fit genotypes to begin to decline, making them appear unfit. Flooding the flask with ancestral λ and running competitions for a short time period (4 hours) solves this problem.

Having a disproportionate number of ancestral λ has a pitfall. We measured λ fitness by comparing the frequency changes of each genotype using amplicon sequencing (more information in the next section). The problem is that most of our sequencing effort would be spent on sequencing a single genotype; the ancestor. To avoid this, we engineered an ancestral genome with 6 synonymous mutations where the reverse primer binds. These edits interfered with primer binding and caused selective amplification of the library λs. We edited 6 'wobble' positions; 3381 (g→t), 3384 (c→a), 3387 (c→a), 3390 (c→t) 3393 (g→a), and 3396 (c→a). The engineering was done using MAGE with the oligo provided in Supplementary Table 10. These edits were made at the end of the protein (3,399 nucleotide) and so they likely did not have an effect on λ fitness. We did not test this because this engineered strain acted as a standard competitor for all competition experiments and so it should have no effect on the relative fitness comparisons.

We ran eight competitions, four in the presence of ancestral *E. coli* and four with *malT*$^{-}$ *E. coli.* Competitions were inoculated with ~$10^7$ total λ particles and ~$2 \times 10^8$ cells into 10 ml of M9 glucose in 50 ml flasks. The cells were preconditioned in M9 glucose for 24 hours before the competition. Flasks were cultured for four hours at 37°C and shaking at 120 rpm. 1 ml samples were removed from the library before the competitions, and after the competition for processing. Phage particles were concentrated using PEG precipitation[11]. The pellet was resuspended in 25 μl of molecular grade water and then a two-step PCR reaction was performed in order to amplify just the region of interest, to attach barcodes to the amplicons so that we could multiplex samples into a single Illumina run, and attach adapters required for Illumina sequencing. The protocol was published by Kelsic *et al.*[13]. The primers we used are provided in Supplementary Table 11. As described in Kelsic *et al.*, we ran two separate PCR reactions for each sample and used a unique barcode on each. This allowed us to test for amplification bias, which we did not detect (Supplementary Fig. 2).

Sequencing

Amplicon samples were pooled together and sequenced using an Illumina Mi-Seq maintained in the Systems Biology Department at Harvard Medical School. 100 base paired end reads were run. We were able to extract on average 107,839 high quality reads per sample with standard deviation of 26,889. This provided considerable coverage to reliably estimate fitness for even rare genotypes.

Post-sequencing analysis and construction of fitness landscape

Amplicon sequences were analyzed to calculate selection rates for each engineered genotype. These values were later used to construct the fitness landscapes. We used a combination of custom Python and MATLAB (version R2019b) scripts to read and concatenate the raw paired-end reads, identify the genotypes based on the focal mutations they possessed, and to count their abundances. All focal mutations were called only if either of the two corresponding watermark mutations were also present. We removed any reads that contained more than one mutation other than the focal and watermark mutations. This essentially acted as a quality filter and we did not have to filter based on the Illumina provided Q-score. For each competition, a mean of counts corresponding to the two barcodes used in PCR amplifications was taken for all the genotypes. If the counts for both, initial and final timepoints were available for a genotype, they were used to estimate the genotype's Malthusian growth rate, given by the calculation $\frac{\ln\frac{\lambda_{i,T}}{\lambda_{i,0}}}{T}$ where $\lambda_{i,t}$ is the density of the given genotype at time $t$ and T is the total time of the competition. Note that most genotypes performed poorly on *malT*— host and fell below detection limit at the final timepoint. We quantified the differences in fitness of all genotypes by calculating their selection rate with respect to the wild type λ (see 'Calculation of selection rate using Malthusian parameter' section). The final fitness landscapes (Fig. 1a and 1b) were constructed by taking the mean of the selection rates obtained from the four replicate competition trials performed on the same host.

Statistical analysis of fitness landscapes

We used multiple linear regression models to quantify the genetic interactions in the fitness landscapes. We regressed fitness values on each landscape against individual effects of the mutations and the pairwise epistatic interactions between them (genotype-by-genotype or GxG interactions). This allowed us to estimate the contribution of epistasis towards explaining the observed fitness data. The fitness of each genotype in the regression model was thus described by 55 predictor variables (10 main effect terms for the ten mutations + 45 interaction terms):

$$y = \beta_{**} + \sum_{i} \beta_{i*} G_i + \sum_{i<j} \beta_{ij} G_i G_j \qquad (2)$$

where $\beta_{**}$ is the intercept, all other $\beta$s are regression coefficients, contribution of an individual mutation $i$ is described by the term $G_i$, and the effect of pair of mutations $i$ and $j$ are captured by terms $G_i G_j$, where $G_i$ is an indicator variable that is equal to 0 when the mutation $i$ is absent and 1 when present. After fitting the model, we used Benjamini-Hochberg procedure[14] to control for false discovery rates and identify statistically significant terms in the model (see Fig. 1c). All analyses were performed in R version 3.6.1[(15)]. The error in fitness estimates of the genotypes did not depend on the mean fitness value of the genotypes (Supplementary Fig. 8).

To understand how λ's landscape differed between the two hosts, we incorporated the host genotype as a predictor variable in the linear model and regressed the combined fitness data of both landscapes together. This allowed us to test if a mutation significantly interacted with the host (genotype-by-environment or GxE interaction) and whether interaction between pairs of mutations changed with the host genotype (genotype-by-genotype-by-environment or GxGxE interaction). The combined full-factorial model consisted of a total of 111 terms (10 G for individual mutation + 1 E for host + 45 GxG + 10 GxE + 45 GxGxE terms),

$$y = \beta_{***} + \sum_i \beta_{i**} G_i + \beta_{**+} E + \sum_i \beta_{i*+} G_i E + \sum_{i<j} \beta_{ij*} G_i G_j + \sum_{i<j} \beta_{ij+} G_i G_j E \quad (3)$$

where additional $E$ is an indicator variable for host type and other notations follow same scheme as in Eq. (2). Since the fit of a model generally improves as the number of predictor variables increases, we tested for overfitting using Akaike information criterion (AIC)[16]. AIC is a penalized-likelihood criterion which penalizes a model for increasing number of parameters in it. The model with the greatest relative likelihood is considered to be the one with minimum AIC value. We minimized AIC value using the *step* function in R[15] to uncover subsets of predictor variables that have high predictive power. This resulted in a more parsimonious model (77 terms out of a total 111, see Fig. 1d). A complimentary adjusted R-squared analysis came to the same conclusion, $R^2_{adj} = 0.775$ for the reduced model and $R^2_{adj} = 0.769$ for the full-factorial model (higher values indicate more predictive and parsimonious models). After model selection, we used Benjamini-Hochberg procedure to determine which variables were significantly predictive (Fig. 1d). This procedure only controls for false positives and is susceptible to false negatives. In this way, it is conservative for our purposes.

Simulation of λ's evolution on fitness landscapes

To test if the changes in λ's fitness landscape facilitated its evolution to infect via OmpF, we simulated λ's evolution on the two landscapes we measured and recorded whether $OmpF^+$ genotypes arose to a high enough frequency that we would have detected them in the original laboratory evolution. We initiated λ populations with no mutations and allowed them to evolve and mutate at any of the ten focal sites used to construct the landscapes (Supplementary Table 1). For each treatment (Fig. 2) we simulated 300 separate trials using a modified Wright-Fisher model with discrete generations and a fixed population size.

For each simulation, a new fitness landscape was constructed by assigning fitness values to all the genotypes ($2^{10}$ = 1024 genotypes). This was done in a way to account for error in estimating fitness and the error associated with imputing the values of missing data points. For the genotypes that had empirical fitness data available (Fig. 1a and 1b), we did not simply average the values of the replicate fitness measurements, but instead we performed a bootstrapping protocol in order to account for the error in the fitness estimate. We randomly resampled from the four replicate measurements (with replacement) and then computed the mean of the four. Some genotypes were not present in all four replicates, in these cases we resampled as many times as there were replicates. Genotypes with only one replicate data point were, thus, assigned the same corresponding fitness values in all the runs of the simulations. Next, we imputed fitness of the genotypes that were missing from the empirical landscape. A genotype with a missing fitness value was randomly chosen and assigned the mean of the fitness values of its nearest neighbors (one-mutation away genotypes) present in the landscape. This was iterated until the full genotypic space was complete. Note that the order in which genotypes are chosen can affect the value that is estimated for a given genotype. This is because as the landscape is filled in, each genotype will have more neighbors to draw values from. This means that if a genotype is randomly chosen early, its fitness will be based on fewer neighbors than if it were chosen later, and its value will be slightly different. Since missing genotypes are randomly chosen in each iteration, the order will vary from one simulation to the next. This method introduces an extra source of variation in the simulation runs and captures uncertainty associated with the imputation of fitness values.

After constructing a complete fitness landscape, we evolved a λ population through repeated cycles of reproduction, selection, and mutation. For each generation, reproduction in the population was simulated by a multinomial sampling where the number of trials was equal to the population size $N$ (set to ~ 6.3 x $10^9$ based on Supplementary Fig. 4) and the success probability associated with a genotype $i$ was given by $p_i = n_i w_i / \sum_i n_i w_i$, where $n_i$ is the abundance of the genotype $i$, and $w_i$ is defined as the exponential of selection rate used in fitness landscape. Thus, the probability of $(k_1, k_2, k_3, \ldots k_m)$ offsprings for genotypes $1,2,3, \ldots, m$ would be:

$$\binom{N}{k_1\ k_2\ k_3 \ldots k_m} \prod_{i=1}^{m} \left( \frac{n_i w_i}{\sum_j n_j w_j} \right)^{k_i}$$

To incorporate mutations, all genotypes whose frequencies increased were mutated as per λ's mutation rate (7.7 x $10^{-8}$ per base per replication[17]). Consider a genotype (say $i$) that increased in abundance and let the number of additional individuals produced by this genotype be denoted by $z_i$ (with $z_i = k_i - n_i$). Each of these $z_i$ individuals retained its parent's genotype with a probability $e^{-\mu}$ (assuming a Poisson distribution for mutations). Otherwise, the individual was assigned to a random neighboring (i.e. mutant) genotype. Given λ's mutation rate, the probability of multiple mutations is very small and was ignored here. We simulated this modified Wright-Fisher cycle of reproduction and mutation for 960 generations. It is difficult to know how many generations phages undergo because the evolved phages have a high spontaneous death rate [10] and can also experience other sources of mortality. Given this, we decided to run the simulation for a somewhat arbitrary amount of time, 960 generations which corresponds to two doublings per hour for the 20-day experiment we are trying to replicate Meyer *et al.*[1]. This is likely an overestimate, which is unintuitively conservative for our purposes because more cycles will cause additional evolution and exploration of the fitness landscape, enhancing the possibility of OmpF$^+$ evolution in the negative controls, and reducing our ability to detect treatment differences.

We evaluated whether λ evolved OmpF$^+$ by examining the population for genotypes that had the necessary mutations to be OmpF$^+$. Previous studies revealed that OmpF genotypes must have four mutations[9]. They all possess two specific changes (A3034G) and (G3319A) and a third change that can occur at positions 3320 or 3321. For the 10 mutations we studied, T3321A is the only third mutation that satisfies this requirement, meaning all OmpF genotypes must have three specific changes (A3034G, G3319A and T3321A). Many J mutations satisfy the last requirement [9], so we implemented what we call the '3+1' rule where genotypes are designated as OmpF$^+$ if they have the three necessary J mutations plus any additional mutation (see Supplementary Table 9). If any such genotype crossed the threshold of 5,000 λ particles during the course of a simulation, the λ population in that run was marked to have evolved OmpF-function. We based the threshold value on the detection limit of OmpF$^+$ genotypes in the original laboratory coevolution experiments (~500 pfu/ml, see 'Coevolutionary Replay Experiments' section).

We implemented host switching by first evolving λ on the landscape measured with ancestral *E. coli,* stopping the simulation early and quantifying the frequency of each λ genotype. Next, we would initiate evolution on the *malT*$^-$ for the remainder of the time but starting with genotypes at the frequencies recorded on the ancestral landscape. A total of 11 different coevolution treatments were run by varying how long λ evolved on the ancestral host landscape and *malT*⁻ landscape (Fig. 2). 300 simulation trials were run for each treatment. We calculated error in the simulations in order to detect significant treatment differences by batching the runs into 30 and estimating the frequency of $OmpF^+$ for each subset. This allowed us to calculate a 95% confidence interval using a Student's *t* distribution for the frequency of $OmpF^+$ evolution in each treatment, and then to use an ANOVA coupled with Tukey's multiple comparison test to test for treatment differences.

Phage competition assays

To test whether λ's path to the innovation requires sequential adaptation to host genotypes isolated from different stages of their coevolution, we studied the dynamics of λ evolution in much more depth than previously reported. We focused on a single experimental replicate reported on in Meyer *et al.*[1]; 'D7' that evolved to be $OmpF^+$ on the 12th day. Daily samples of the population were preserved so we were able to revive phages from the full time series. We isolated 66 phages in total (Supplementary Table 3 & Supplementary Table 4) and sequenced the full-length J gene from each in order to reconstruct the evolutionary dynamics (Fig. 3a).

Next, we ran head-to-head competition experiments from key genotypes along the path to $OmpF^+$, the wildtype (WT) and λ with a single *J* mutation (A), double (AB), and quintuple (ABCDE) (Fig. 3b and 3c, Supplementary Fig. 6). Our first step was to mark key strains with a gene that caused the plaques to be visually distinguishable from unmarked phage. The gene is *lacZ*α, which in the presence of a 5-bromo-4-chloro-3-indolyl-b-D-galactopyranoside (X-gal), Isopropyl-β-D-thiogalactoside (IPTG), and a host cell like DH5α that lacks *lacZ*α, produces blue plaques. To incorporate *lacZ*α into λ's genome, we used a gene fusion of *lacZ*α with λ's *R* gene[18,19]. The fusion readily recombines into λ's genome by the phage's endogenous recombination system, λ-red[20]. The procedure is straightforward, infect an *E. coli* strain that has a plasmid with the lacZ fusion (strain: SYP042; plasmid: pSwtRlaczalphaZalpha+RZ xl1 Blue Amp Blue provided by Ing-Nang Wang, Albany, NY) and some fraction of the λ produced will have recombined with the plasmid. The recombinants are isolated by picking blue plaques. WT and AB were each marked. The marker has been shown to have a slight fitness effect, however, this would not significantly influence our measurements of fitness since we observed such large differences between strains[18,19].

We competed three pairs of phages, $WT_{lacZ}$ versus A, A versus $AB_{lacZ}$, and $AB_{lacZ}$ versus ABCDE, under two conditions: with ancestral *E. coli* or *malT*⁻. Competitions were run for a single 24-hour period under identical conditions in which the phage evolved[1]. Initial λ density was between $10^4$-$10^5$ particles per ml and the relative frequency of the strains was sometimes skewed in order to start with more of the unfit genotype. Fitness differences were so large that the less fit genotype would be overwhelmed and its frequency undetectable if they did not start with a numerical advantage. ~$5x10^6$ exponentially growing cells were inoculated to each flask ($5x10^5$ per ml). The cells were preconditioned by growing them overnight in the competition medium. Three replicate competitions were run for each treatment. Phage densities of the unmarked and marked λ were determined by plating in typical soft agar plates, where we added 0.5 mg/ml of X-gal and 0.25 mg/ml IPTG to the molten soft agar. Densities were measured at t=0 and t=24 h.

Host competition assay

This competition experiment tested the role of λ's evolution plays in promoting the evolution of host resistance. We competed the ancestral host (REL606) against the evolved-resistant host (EcC4) in the presence of different λ genotypes that had increasing numbers of *J* mutations (Fig. 3; WT (cI26), A (1-mutation), ABC (3-mutation) and A**C (4-mutation)). We initiated 3 replicate populations for each phage treatment. The competition was performed with identical conditions as the original evolution experiment and run for 4 h to prevent λ from evolving during the competition assay. Cells were preconditioned in modified M9 glucose for 24 hours before the competition. As with λ, fitness was measured by observing the change in frequency of the competitors over time. Frequencies were determined by plating a subsample of the populations onto tetrazolium maltose (TMal) agar plates[21]. REL606 produces white colonies on the plates, while EcC4 produces smaller red colonies. Before plating, the phages were removed by centrifugation of the cells and then removing the supernatant that possessed the phage. Cells were resuspended in saline solution and the centrifugation and resuspension was repeated once more.

<u>Coevolutionary replay experiments</u>

Would λ's evolution to OmpF innovation proceed without initially evolving on ancestral host? We tested this by replaying λ-*E. coli* coevolution but starting out with a host that already evolved resistance via a *malT*⁻ mutation (strain LR01). We initiated 12 replicate populations with *malT*⁻, and another 12 with REL606 as a positive control. The replay experiment was run nearly identically to the coevolution experiment performed by Meyer *et al.*[1] with cI26 as the ancestor phage, and daily sampling was done to detect presence of $OmpF^+$ phage by spotting the phage on bacterial lawn of *lamB*⁻ cells (detection limit ~500 pfu per ml) (Supplementary Fig. 7). The two differences were that the study was run longer (31 days) and daily estimates of λ populations were made (Supplementary Fig. 4). 12 replicates were chosen in order to have enough statistical power to determine whether or not a treatment reduces the chances of $OmpF^+$ evolution. In the previously reported study, only a quarter of populations (24 out of 96) evolved to use OmpF[1]; this result has also been replicated in our lab on two separate instances where 3 out of 12 populations gained OmpF-function. Assuming a binomial distribution and a *true* success rate of 0.25 for λ's $OmpF^+$ evolution with ancestral host, the probability of observing no replicate evolving to be $OmpF^+$ in 12 replicate populations is 0.03167. Thus, we can conclude that *malT*⁻ treatment reduced λ's ability to evolve the innovation as compared to when coevolution is initiated with ancestral host. Two isolates from each population were sampled at day 26 and the reactive region of their *J* genes were sequenced using the Sanger sequencing method.

We ran two additional replay experiments as a positive control for any unintended side effects of initiating an experiment with *malT*⁻ host (LR01). A prediction from the fitness landscape simulation results was that if we started a replay with *malT*⁻ and phage isolated from a later-stage of the coevolution, then the phage should evolve OmpF use. To test this, we genetically modified an $OmpF^+$ λ by removing one mutation required for OmpF use ($\lambda^{-1}$), and then a second ($\lambda^{-2}$). The $OmpF^+$ genotype was a lysogen (cI857) that we had previously edited in 7 *J* mutations (Supplementary Table 6), which is reported about in Petrie *et al.*[10]. The two new edits were made using MAGE and the oligos are reported in Supplementary Table 10.

The replay experiments were run identically, except because of an oversight, these replays were run in a slightly different minimal glucose medium called Tris DM (compared to original coevolution experiment run by Meyer *et al.*[1]). The key difference in the medium is the buffer, it is a Tris buffer, not a phosphate buffer. Both mediums impose carbon limitation and have the same concentration of the single carbon source, glucose, and so population and evolutionary dynamics should not have been affected. We confirmed in an additional replay experiment that the medium does not affect the timing or repeatability of $OmpF^+$ evolution.

#### Supplementary Discussion

Fitness effect of synonymous mutations used as watermarks in MAGE

We designed two watermark mutations for each focal mutation. Watermark mutations are synonymous mutations that fall within a few nucleotides of the focal mutation. These edits improved our ability to detect the presence of focal mutations within the MAGE library (Supplementary Table 9). In principle, these watermark mutations should not affect the fitness of λ because the introduced mutations do not change the amino acids. However, we designed two different watermarks in order to test whether or not they influence λ fitness. To test for fitness effects of the synonymous mutations, we evaluated the fitness of genotypes with just a single focal mutation, and then we split the count data from the competition experiments into two, one for watermark '1' and a second pool for '2'. We calculated the fitness of each group and compared their means using a *t*-test. The analysis was limited to just fitness measurements made for the treatment with the ancestral host because the single mutants did poorly on *malT*$^{-}$ host and many dropped below our limit of detection during the competitions. We were also only able to run the analysis on 7 out of 10 mutations because in three cases only one watermark was represented in the final counts data. We found that the fitness did not depend on the neutral marker for all seven using a Bonferroni corrected alpha value of 0.0071 (Table 12). However, one of the seven was significant based on the uncorrected alpha value of 0.05. The effect on the selection rate was estimated to be 0.9947 with 95% confidence interval of 0.4758 to 1.5136. This value falls well within the error associated with fitness estimates (Supplementary Fig. 8). Because of our uncertainty of whether there was an effect of this synonymous mutation, and because the effect falls within normal levels of error, we did not take steps to correct for this possible effect.

Simulation results when only using genotypes present in both the ancestral and *malT*$^{-}$ fitness landscapes

One possible complication with the fitness landscape analyses is that the *malT*$^{-}$ landscape is based on a subset of the ancestral landscape (131 versus 580 out of 1,024 possible genotypes). This could lead to potentially spurious comparisons of evolution on the two hosts. To control for this, we re-ran the simulations starting with only the genotypes that were present in both the landscapes. While the frequencies of $OmpF^+$ populations did shift with this new analysis, the main result that switching landscapes enhances the frequency of $OmpF^+$ evolution remained statistically significant (Supplementary Fig. 5d). The increased frequency in ancestor-only treatment stems from the increased stochasticity in estimating the landscape since more genotypes are imputed in panel d than a-c. This stochasticity leads to more $OmpF^+$ evolution because the increased randomness in landscape formation increases the chances of producing viable pathways to OmpF use. Interestingly, the ancestral landscape $OmpF^+$ frequency is equivalent to *malT*$^{-}$ in panel d. This similarity suggests that the frequency of OmpF evolution for *malT*$^{-}$ may be artificially high, which would mean that in reality there may be greater differences between this treatment and the fluctuating treatments.

Simulation results when shifting is done between landscapes of the same host

Each time we impute fitness values of missing genotypes in a simulation run, a slightly different landscape is produced. This procedure raises the question of whether the increased frequency of $OmpF^+$ in the shifting landscape simulations is due to structural differences between the two landscapes, or the random differences created by the imputation technique. To test this, we repeated simulations as previously discussed; however, this time we shifted to a newly generated landscape created from the same host-landscape data. $OmpF^+$ did *not* evolve when the shift was made between two ancestral landscapes. λ did evolve OmpF function in some replicates when the switching was made between two *malT*— landscapes; however, the frequency was significantly less than when the shift was made from ancestor to *malT*— (Supplementary Fig. 5e and 5f). Lastly, to control for the sparser landscape of *malT*— compared with the ancestor, we shifted between two ancestral host landscapes that were generated using only fitness values of genotypes present in both landscapes. We found the same qualitative result as for the *malT*— to *malT*— landscape shift (Supplementary Fig. 5g and 5h). These results show that the structural differences between the two host landscapes encourage $OmpF^+$ evolution above and beyond what results from the noise associated with the imputation procedure.

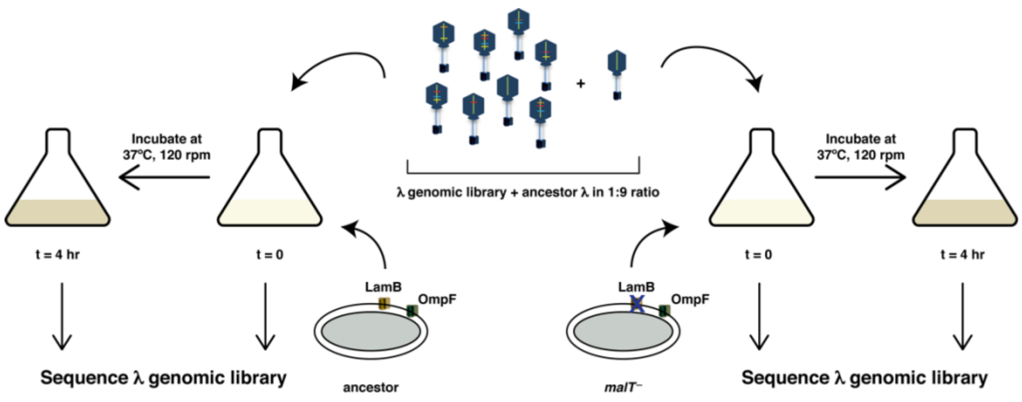


**Supplementary Fig. 1. Schematic illustration showing how fitness was measured for the combinatorial λ library in order to construct fitness landscapes.** The genomic library was first mixed with the ancestor in 1:9 ratio, incubated with different types of hosts for four hours at 37°C, and then sequenced at both initial and final timepoints to calculate the selection rates of individual genotypes.

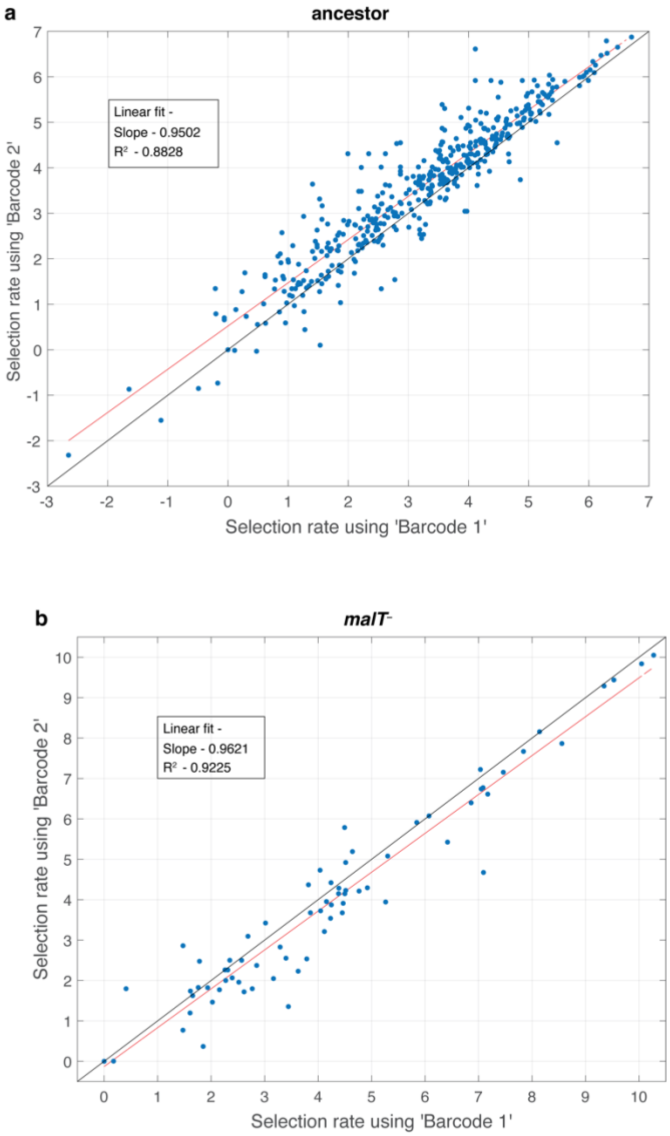


**Supplementary Fig. 2. Test to determine whether amplification distorts measurements of fitness.** For every replicate population in the competition (total eight, four for each host) we performed two separate amplifications and used a unique barcode for each. This allowed us to compare fitness calculations based on independent amplifications and to determine if the amplification step is repeatable and unbiased. Each point represents a unique λ genotype presented in the fitness landscapes reported in Fig. 1. The red line is a linear least-square fit. The regression fitting parameter and coefficient of determination are provided in the inset. The black line represents the vector where fitness estimates from each barcode are equal.

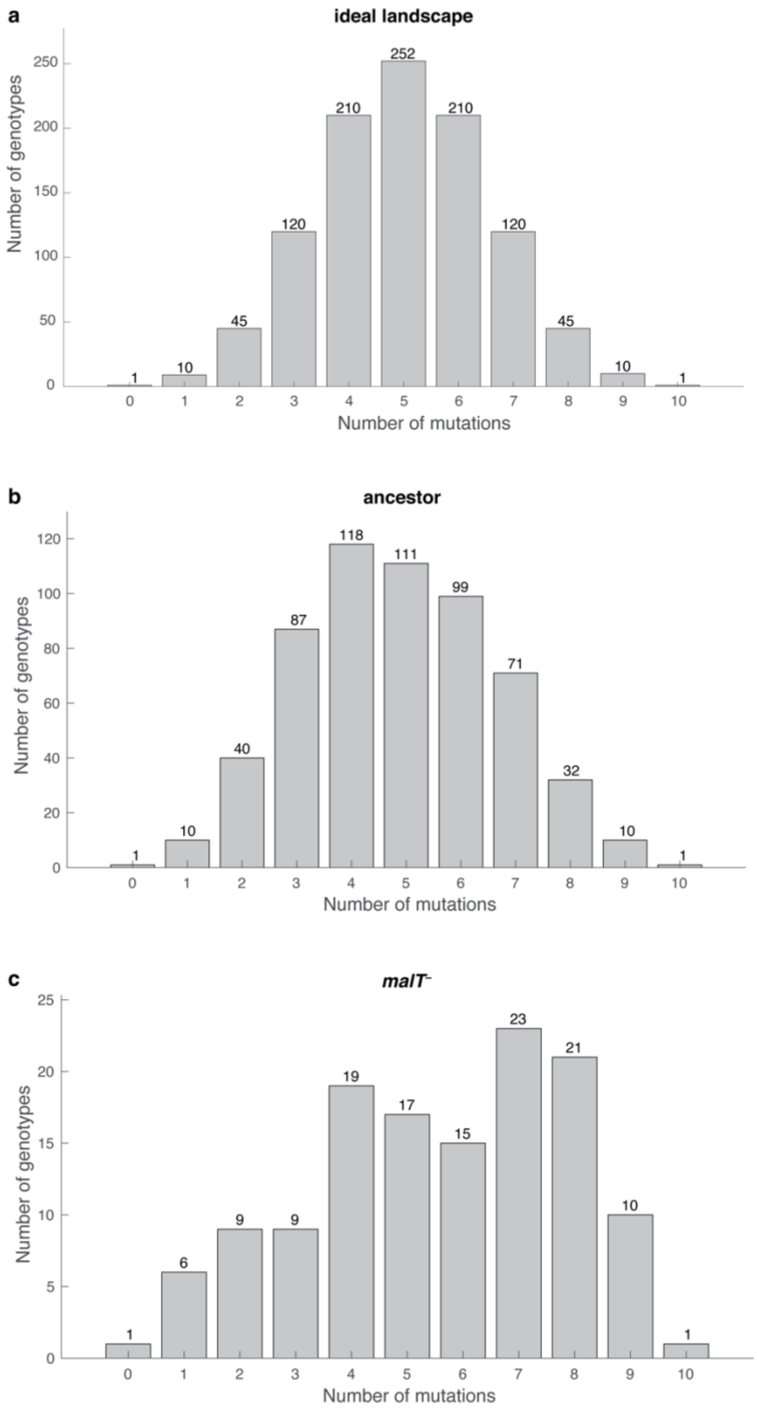


**Supplementary Fig. 3. Distribution of λ genotypes present with respect to the number of mutations they possess. a)** An ideal fitness landscape where all combinations of ten mutations are present. **b) and c)** Empirical fitness landscapes with ancestral host and *malT*— host, respectively.

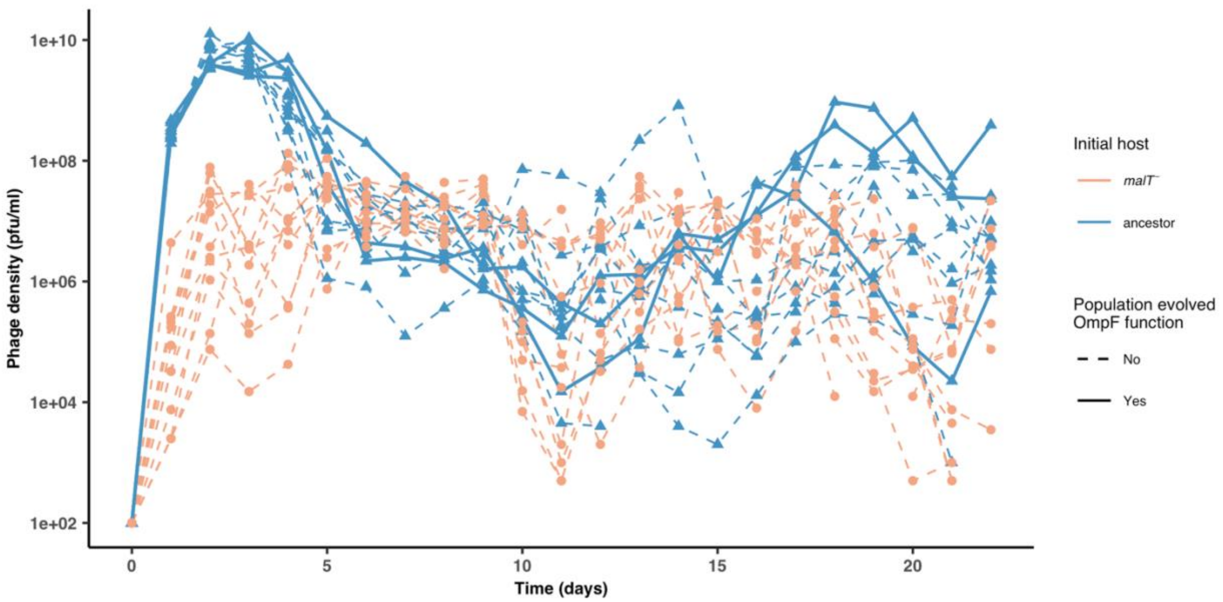


**Supplementary Fig. 4. Phage densities during the coevolutionary replay experiment performed in Fig. 4.** Timeseries of phage density of 12 replicates measured at the end of each day for cultures initiated with wildtype λ and ancestor host or *malT*— host (corresponding to Fig. 4a and 4b). The three λ populations that evolved OmpF function in the ancestor host treatment are marked by solid lines. For the simulations, we used a populations size of 6.3 x $10^9$ which is lower than the peak population density observed (~$10^{10}$), but above the average population density (~ 6.3 x $10^8$).

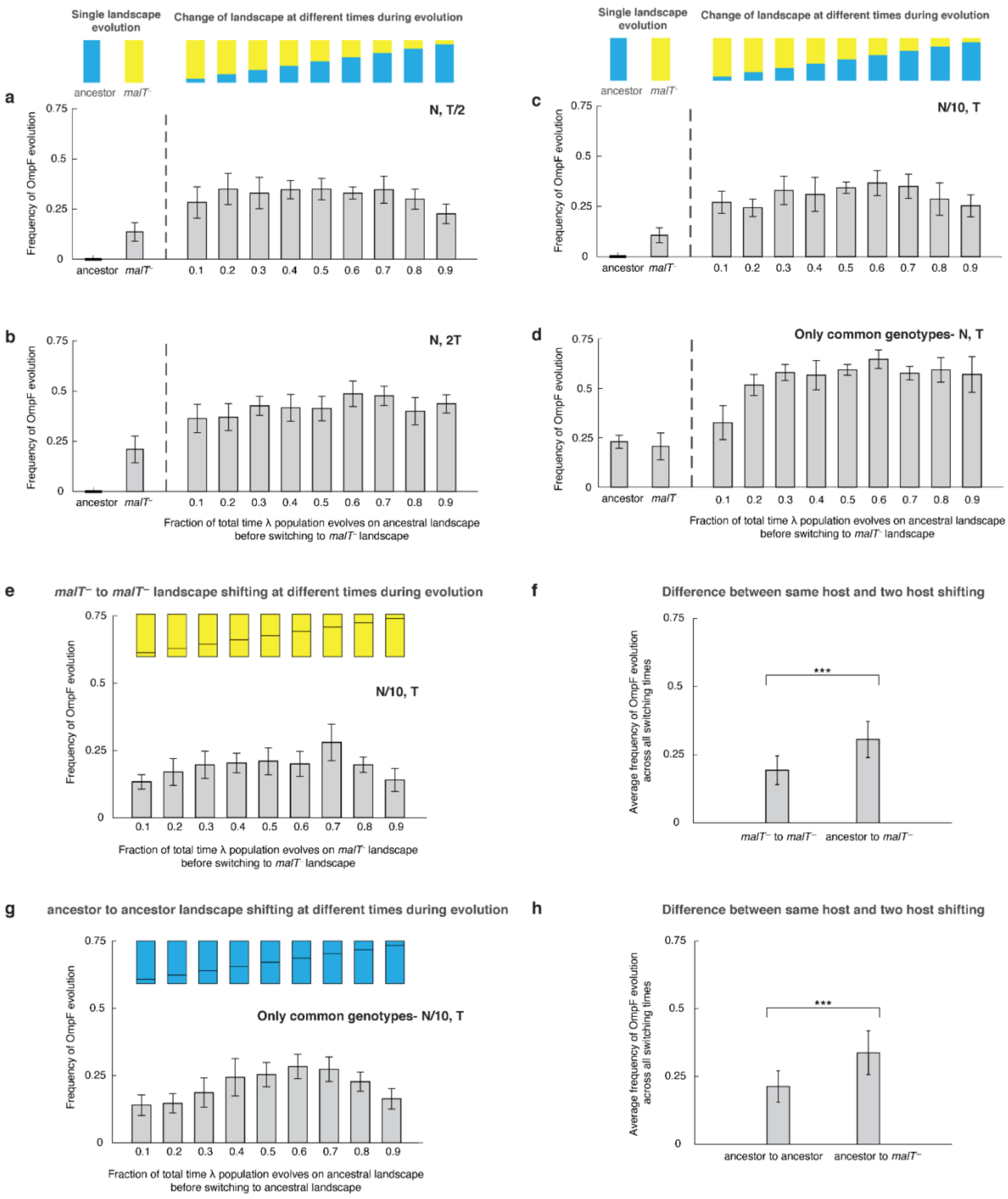


**Supplementary Fig. 5. Additional simulations to verify that shifting landscapes promote OmpF$^+$ evolution.** Frequency of OmpF$^+$ evolution when λ population is evolved on different landscapes with different controls as compared with Fig. 2— a) half the total number of generations, b) twice the total number of generations, c) with one-tenth of the population size, and d) starting with only the genotypes whose fitness values were measured for both the landscapes. For all switching landscape treatments except two (switching fraction-time of 0.9 in (a), and 0.1 in (d)), the frequency of $\text{OmpF}^+$ evolution was significantly higher (adjusted P-value<0.05) than the constant landscape treatments (ANOVA: $F$-ratio = 6.62, $d.f. = 99$, $P$ < 0.0001 for (a), $F$-ratio = 8.22, $d.f. = 99$, $P$ < 0.0001 for (b), and $F$-ratio = 8.12, $d.f. = 99$, $P$ < 0.0001 for (c), and $F$-ratio = 36.04, $d.f. = 109$, $P$ < 0.0001 for (d)). Each bar represents an average of 300 simulation runs and error bars indicate 95% confidence interval. The frequency of $\text{OmpF}^+$ evolution in our simulations depends mainly on whether λ spends any time on the ancestral landscape before shifting to $malT^-$ and there is little evidence of a relationship between the amount of time spend on each landscape and the probability of evolving $\text{OmpF}^+$. However, we can reason that as the time spent on the ancestral landscape approaches zero, the frequency of $\text{OmpF}^+$ will drop and approach the frequency caused by the constant $malT^-$ landscape. This transition must happen sometime between 0 and 10%. e) and g) show the frequency of $\text{OmpF}^+$ evolution when shifting is done between the same hosts, and f) and h) show difference between this switching treatment, and the varying host treatment. The bars are the averages taken across all switching times. Statistics for two-sample t-tests: f) $t = 9.10, d.f. = 169, P < 0.0001$; h) $t = 8.54, d.f. = 161, P < 0.0001$.

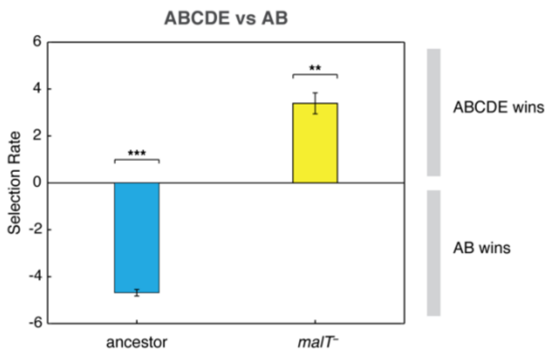


**Supplementary Fig. 6. Competition assay of λ isolates from population D7.** Relative fitness of λ with two mutations (AB) to an OmpF$^+$ λ with five mutations (ABCDE) on two host genotypes. This shows that AB outcompetes ABCDE in the presence of ancestral *E. coli* while ABCDE is favored over AB with *malT*$^-$ *E. coli.* Selection rate (per 4 h) is the difference in Malthusian growth rates of the competitors over one day with a value of 0 indicating no difference in fitness. Each bar represents mean of three replicate trials and error bars indicate one sample standard deviation. (One-tailed t-test to test if the selection rate is greater (or less) than zero— ABCDE vs AB with ancestor host: $t = -58.53, d.f. = 2, P < 0.0001$, ABCDE vs AB with *malT*$^{-}$: $t = 13.09, d.f. = 2, P = 0.0029$. Asterisks over the bar graphs indicate significance level corresponding to the P-values).

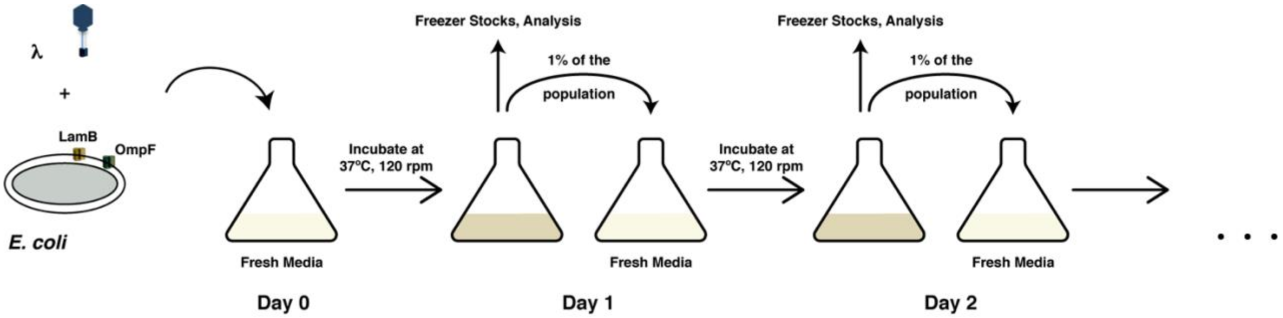


**Supplementary Fig. 7. A schematic overview of the coevolution experiments performed in this study and by Meyer *et al.*[1].** Coevolution is initiated by coculturing λ and *E. coli* in glucose media for 24 hours at 37°C. At the end of each day, 1% of the total population is transferred to fresh media to continue the coevolution and the spent media is discarded. A fraction of the community is also sampled for long-term storage at -80°C, to determine population densities and to check the status of λ's OmpF use.

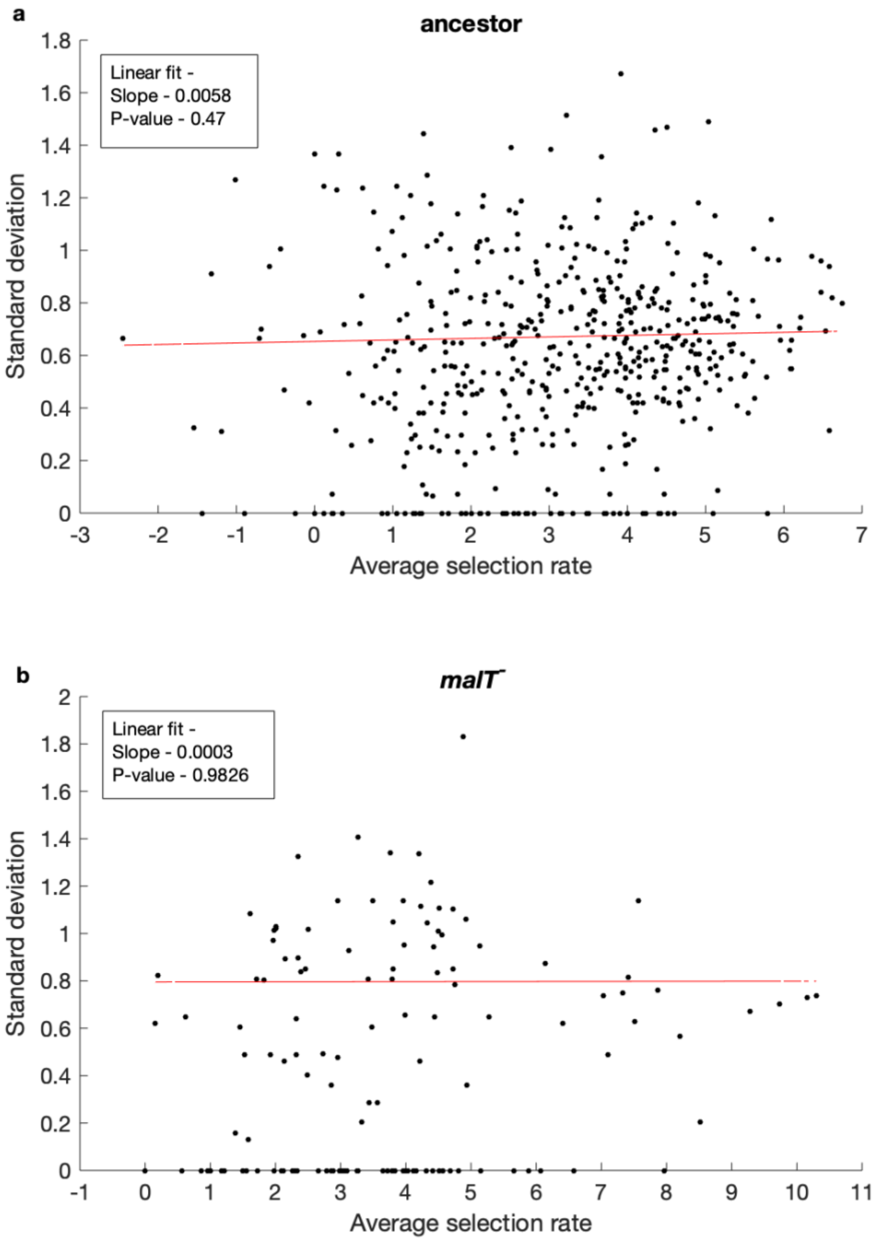


**Supplementary Fig. 8. Test of whether the error in our estimates of selection rates (per 4 h) is influenced by the magnitude of the estimate.** Standard deviation does not correlate with mean fitness of a genotypes for both a) ancestor host, and b) *malT*⁻ host; P-values in the insets indicate that the slope is not significantly different than 0. Uniform variance in fitness values permit the performance of analysis of variance to detect genotype by genotype by host interactions. Points with zero standard deviation indicate presence of only one replicate population with a fitness value for that genotype.

**Supplementary Table 1.** Reconstruction of the list of mutations in 24 phage isolates that evolved to target the OmpF receptor in the large-scale coevolution experiment by Meyer *et al.*[1]. Highlighted mutations were chosen to form the genotype space for the 10-dimensional fitness landscape in Fig. 1. The first five and last five mutations fall within two 100-nucleotide windows for Mi-Seq 100 base paired-end sequencing.

| λ genotypes which evolved OmpF+ function in (*1*) / **Mutations in *J* →** | T487C | C494T | C599T | A1747G | C2879T | A2966G | C2969T (red) | A2975C | C2988A | A2989G (red) | T2991G (red) | C2999T (red) | C3033T | A3034G (red) | C3119T | T3143A | C3147G | A3158A | C3227T | T3230C | A3233G | T3248C | C3310T (red) | G3319A (red) | A3320G (red) | T3321G | T3321A (red) | A3364T | T3380C (red) |
|---|---|---|---|---|---|---|---|---|---|---|---|---|---|---|---|---|---|---|---|---|---|---|---|---|---|---|---|---|---|
| A7 | | | | | | | | | | ■ | | ■ | | ■ | ■ | | | | | | | | ■ | ■ | | | ■ | | |
| A8 | | | | | | | | | | ■ | | ■ | | ■ | ■ | | | | | | | | ■ | ■ | | | ■ | | |
| A12 | | | ■ | | | ■ | | | | | ■ | | ■ | ■ | | ■ | ■ | | | | | | | ■ | | | ■ | | ■ |
| B2 | | | | | | | ■ | | | | | | | ■ | ■ | | | | | | | | | ■ | | | ■ | | |
| C2 | | | | | | | | | | ■ | | ■ | | ■ | ■ | | | | | | | | ■ | ■ | | | ■ | | |
| C3 | | | | | | | | | ■ | | | | | ■ | | | | | | ■ | | | | ■ | | | ■ | | |
| D3 | | | | | | | | | | | ■ | | | ■ | ■ | | | | | | | | ■ | ■ | | | ■ | ■ | |
| D4 | | | | | | | | | | | | ■ | | ■ | | | | | ■ | | | | | ■ | | | ■ | | ■ |
| D6 | ■ | ■ | | | | | ■ | | | | | | | ■ | ■ | | | | | | | | | ■ | | | ■ | | |
| D7 | | | | | | | | | | | ■ | | | ■ | | | | | | | | | | ■ | | | ■ | | |
| D9 | | | | | ■ | | | | | | ■ | | | ■ | ■ | | | | | | | | | ■ | | | ■ | | |
| E3 | | | | | | | ■ | | | | | ■ | | ■ | | | | | | ■ | | | ■ | ■ | | | ■ | | |
| E4 | | | | | ■ | | | | | | | ■ | | ■ | ■ | | | | | | | ■ | ■ | ■ | ■ | | | | |
| E11 | | | | | | | | | | ■ | | ■ | | ■ | ■ | | | | | | | | ■ | ■ | | | ■ | | |
| E12 | | | | | | | | | | ■ | | ■ | | ■ | ■ | | | | | | | | ■ | ■ | | | ■ | | |
| F5 | | | | | | | | ■ | | | | ■ | | ■ | | | | ■ | | ■ | | | | ■ | | | ■ | | |
| F7 | | | | | ■ | | | | | | ■ | | | ■ | ■ | | | | | | | | | ■ | | | ■ | | |
| F8 | | | | | ■ | | ■ | | | | | ■ | | ■ | | | | | | ■ | | | ■ | ■ | | | ■ | | |
| G4 | | | | | | | ■ | | | | | | | ■ | ■ | | | | | | | | | ■ | | | ■ | | |
| G9 | | | | ■ | | | | | | ■ | | ■ | | ■ | ■ | | | | | | | | | ■ | | ■ | | | |
| H5 | | | | | | | | | | ■ | | ■ | | ■ | ■ | | | | | | | | ■ | ■ | | | ■ | | |
| H8 | | | | | | | | | | ■ | | ■ | | ■ | ■ | | | | | | | | ■ | ■ | | | ■ | | |
| H9 | | | | | ■ | | | | | | ■ | | | ■ | ■ | | | | | | | | | ■ | | | ■ | | |
| H12 | | | | | | | | | ■ | | | | | ■ | | | | | | ■ | | | ■ | ■ | | | ■ | | |

**Supplementary Table 2.** Result of Tukey's significant test comparing for difference of means between different types of simulations in Fig 1c. The simulation treatments that were significantly different from each other are marked in red.

| Difference of levels | Adjusted P-value | Difference of levels | Adjusted P-value | Difference of levels | Adjusted P-value |
|---|---|---|---|---|---|
| $malT^{-}$ - 0.1 | 0.0124 | 0.1 - 0.8 | 0.9907 | 0.4 - 0.5 | 0.9838 |
| $malT^{-}$ - 0.2 | 0.2657 | 0.1 - 0.9 | 0.9585 | 0.4 - 0.6 | 0.1295 |
| $malT^{-}$ - 0.3 | 0.0003 | 0.2 - 0.3 | 0.4122 | 0.4 - 0.7 | 0.3106 |
| $malT^{-}$ - 0.4 | 0.0691 | 0.2 - 0.4 | 0.9999 | 0.4 - 0.8 | 0.8427 |
| $malT^{-}$ - 0.5 | 0.0023 | 0.2 - 0.5 | 0.7988 | 0.4 - 0.9 | 0.6969 |
| $malT^{-}$ - 0.6 | <0.0001 | 0.2 - 0.6 | 0.0269 | 0.5 - 0.6 | 0.7499 |
| $malT^{-}$ - 0.7 | <0.0001 | 0.2 - 0.7 | 0.0859 | 0.5 - 0.7 | 0.9384 |
| $malT^{-}$ - 0.8 | 0.0004 | 0.2 - 0.8 | 0.4676 | 0.5 - 0.8 | 1.0000 |
| $malT^{-}$ - 0.9 | 0.0002 | 0.2 - 0.9 | 0.3106 | 0.5 - 0.9 | 0.9989 |
| 0.1 - 0.2 | 0.9733 | 0.3 - 0.4 | 0.7988 | 0.6 - 0.7 | 1.0000 |
| 0.1 - 0.3 | 0.9838 | 0.3 - 0.5 | 0.9999 | 0.6 - 0.8 | 0.9585 |
| 0.1 - 0.4 | 0.9999 | 0.3 - 0.6 | 0.9733 | 0.6 - 0.9 | 0.9907 |
| 0.1 - 0.5 | 1.0000 | 0.3 - 0.7 | 0.9989 | 0.7 - 0.8 | 0.9976 |
| 0.1 - 0.6 | 0.4122 | 0.3 - 0.8 | 1.0000 | 0.7 - 0.9 | 0.9999 |
| 0.1 - 0.7 | 0.6969 | 0.3 - 0.9 | 1.0000 | 0.8 – 0.9 | 1.0000 |

**Supplementary Table 3.** Mutations and their corresponding labels in λ genotypes isolated from population D7 in Meyer *et al.*[1]. Red asterisks indicate the particular mutation in a genotype's description in Fig. 3a and Supplementary Table 4.

| Mutations / λ Isolates | T2991G (A) | A3031G (A*) | A3034G (E) | G3319A (B) | A3320G (A**) | T3321A (D) | T3380C (C) |
|---|---|---|---|---|---|---|---|
| WT | | | | | | | |
| A | ■ | | | | | | |
| A* | ■ | ■ | | | | | |
| A** | ■ | ■ | | | ■ | | |
| AB | ■ | | | ■ | | | |
| ABC | ■ | | | ■ | | | ■ |
| ABCDE | ■ | | ■ | ■ | | ■ | ■ |

**Supplementary Table 4.** Relative abundance of λ genotypes present at different times in population D7. This data was used to create the Muller plot in Fig. 3a.

| Day | Total number of plaques picked | Genotype | Frequency of the genotype |
|---|---|---|---|
| 0 | - | WT | 1 |
| 1 | 1 | A | 1 |
| 2 | 1 | A | 1 |
| 3 | 1 | A | 1 |
| 4 | 5 | A | 1 |
| 5 | 5 | A | 1 |
| 6 | 11 | A | 0.273 |
| | | A** | 0.636 |
| | | A* | 0.091 |
| 7 | 11 | A** | 0.818 |
| | | ABC | 0.091 |
| | | A**C | 0.091 |
| 8 | 11 | ABC | 0.273 |
| | | A**C | 0.727 |
| 9 | 5 | ABC | 0.2 |
| | | A**C | 0.8 |
| 10 | 5 | ABC | 1 |
| 11 | 5 | A**C | 0.2 |
| | | ABC | 0.2 |
| | | ABCDE | 0.6 |
| 12 | 5 | ABCDE | 1 |

**Supplementary Table 5.** Mutations present in λ isolates from day 26 of the coevolutionary replay experiment initiated with a) ancestor host and b) *malT*⁻ host (corresponding to Fig. 4a and 4b). Two strains (*a* and *b*) were isolated from each population and the active region of *J* (roughly between nucleotide position 2,600 and the end) sequenced. Replicates marked in red indicate populations that evolved OmpF-function. Canonical mutations for evolution of OmpF function are bolded. λ evolved more of both the total number of mutations and total number of canonical mutations in replicate populations initiated with ancestor host than with the evolved *malT*⁻ host (statistics for difference in total number of mutations: $t = 5.37, d.f. = 11, P = 0.0002$, statistics for difference in total number of canonical mutations: $t = 3.18, d.f. = 11, P = 0.0088$; both were tested using two-sample t-test with unequal variances assumed).

| | **Mutations in *J*** → | A2866T | T2908A | G2921A | G2966T | C2969T | C2988A | C2988G | A2989G | T2991G | T2993C | C2999T | A3031G | **A3034G** | C3119T | C3147G | G3226T | C3227T | T3230C | C3310T | **G3319A** | **T3321G** | **T3321A** | T3331C | **Total number of mutations** |
|---|---|---|---|---|---|---|---|---|---|---|---|---|---|---|---|---|---|---|---|---|---|---|---|---|---|
| **ancestor host** | 3a | | | | | | ■ | | | | | ■ | | | | | | | | | ■ | | | | 3 |
| | 3b | | | | | | ■ | | | | | ■ | | | | | | | | | ■ | | | | 3 |
| | 4a | ■ | | | | | | | | | | ■ | | ■ | ■ | | | ■ | | ■ | ■ | | ■ | | 8 |
| | 4b | ■ | | | | | | | | | | ■ | | ■ | ■ | | | ■ | | ■ | ■ | | ■ | | 8 |
| | 5a | | | ■ | | | ■ | | | | | | | | | ■ | | ■ | | | | | | | 4 |
| | 5b | | | ■ | | | ■ | | | | | | | | | ■ | | ■ | | | | | | | 4 |
| | 6a | | | | | | | | | ■ | ■ | | ■ | | ■ | | | | | | ■ | | | | 5 |
| | 6b | | | | | | | | | ■ | ■ | | ■ | | ■ | | | | | | ■ | | | | 5 |
| | 8a | | | | | | | | | | | | | ■ | ■ | | | ■ | | | ■ | | | | 4 |
| | 8b | | | | | | | | | | | | | | ■ | | | ■ | | | ■ | | | | 3 |
| | 9a | | | | | | | | ■ | ■ | | | | | ■ | | | | | | ■ | | | | 4 |
| | 9b | | | | | | | | ■ | ■ | | | | | ■ | | | | | | ■ | | | | 4 |
| | 10a | | ■ | | | | | | | ■ | | | | | ■ | | | | | | ■ | | | | 4 |
| | 10b | | ■ | | | | | | | ■ | | | | | ■ | | | | | | ■ | | | | 4 |
| | 11a | | | | | | ■ | | | | | | | ■ | ■ | | | | | ■ | ■ | ■ | | | 6 |
| | 11b | | | | | | ■ | | | | | | | ■ | ■ | | | | | ■ | ■ | ■ | | | 6 |
| | 12a | | | | | | | ■ | | | | ■ | | ■ | ■ | | | | | | ■ | ■ | | | 6 |
| | 12b | | | | | | | ■ | | | | ■ | | ■ | ■ | | | | | | ■ | ■ | | | 6 |
| ***malT*⁻ host** | 2a | | | | | | | | | | | | | | ■ | | | | | | | | | | 1 |
| | 2b | | | | | | | | | | | | | | ■ | | | | | | | | | | 1 |
| | 4a | | | | | ■ | | | | | | | | | | | | ■ | | | | | | | 2 |
| | 4b | | | | | | | | | | | | | | | | | ■ | | | | | | | 1 |
| | 5a | | | | ■ | | | | | | | | | | | | | | | | | | | ■ | 2 |
| | 5b | | | | ■ | | | | | | | | | | | | ■ | | | | | | | ■ | 3 |
| | 6a | | | | | | | | | | | | | | | | | | ■ | | | | | | 1 |
| | 6b | | | | | | | | | | | | | | | | | | ■ | | | | | | 1 |
| | 8a | | | | | | | | | ■ | | | | | | | | | | | ■ | | | | 2 |
| | 8b | | | | | | | | | ■ | | | | | | | | | | | ■ | | | | 2 |

**Supplementary Table 6.** List of mutations in 7-mut OmpF$^+$ cI857 lysogen and the two engineered OmpF$^—$ genotypes; - $\lambda^{-1}$ and $\lambda^{-2}$ (see 'Coevolutionary replay experiments' in Methods for details on construction). Bolded mutations represent the three canonical mutations for OmpF$^+$ function.

| **λ genotype** | **Mutations** |
|---|---|
| **7-mut (OmpF$^+$)** | C2999T, **A3034G**, T3230C, C3310T, **G3319A**, **T3321A**, A3364T |
| **$\lambda^{-1}$ (OmpF$^—$)** | C2999T, **A3034G**, T3230C, C3310T, **G3319A**, A3364T |
| **$\lambda^{-2}$ (OmpF$^—$)** | C2999T, T3230C, C3310T, **G3319A**, A3364T |

**Supplementary Table 7.** Pairwise P-value for Tukey's HSD (honestly significant difference) test comparing difference of means between different competitions in Fig. 3d.

| Difference of levels | Adjusted P-value |
|---|---|
| WT - A | 0.0022 |
| WT - ABC | <0.0001 |
| WT - A**C | <0.0001 |
| A - ABC | <0.0001 |
| A - A**C | <0.0001 |
| ABC - A**C | 0.9998 |

**Supplementary Table 8.** PCR primers for sequencing *J* gene in λ.

| | PCR primers |
|---|---|
| **Forward Primer (5'-3')** | CCTGCGGGCGGTTTTGTCATTTA |
| **Reverse Primer (3'-5')** | CGCATCGTTCACCTCTCACT |

**Supplementary Table 9.** List of focal mutations with their corresponding two watermark mutations. The three canonical mutations required for OmpF-function[9] are marked in bold.

| Focal *J* Mutation | Mutation ID used for linear regression analysis | Amino Acid Change | Neutral 1 | Neutral 2 |
|---|---|---|---|---|
| C2969T | G1 | A → V | G2970C | G2970T |
| A2989G | G2 | I → V | G2985T | C2988T |
| T2991G | G3 | I → M | A2994C | A2994G |
| C2999T | G4 | A → V | G3000A | G3000C |
| **A3034G** | G5 | S → G | C3033T | T3036A |
| C3310T | G6 | H → Y | G3309A | G3309T |
| **G3319A** | G7 | D → N | G3315C | G3315A |
| A3320G | G8 | D → G | A3318C | A3318G |
| **T3321A** | G9 | D → E | T3324A | T3324C |
| T3380C | G10 | L → P | G3378A | G3378C |

**Supplementary Table 10.** Oligos used in MAGE to insert *J* mutations in λ.

| **Mutations introduced** | **Sequence (5'-3')** |
|---|---|
| A3321T | CATCGCTGGCAAACGTATACGGCGGAATaTTTGCCGAATACCGTGT GGACGTAAGCGTGAACGTCAGGATCACGTTTCCCCGACCCGCTG |
| G3034A | CATCGGTCACGGTGACAGTACGGGTACCTGACGGCCAGTCCACACt GCTTTCACGCTGGCGCGGAAAAGCCGCGCTCGCCACCTTTACAA |
| 6 ‘wobble’ edits- G3381T C3384A, C3387A, C3390T, G3393A, and C3396A | TAAAACGCCCGTTCCCGGACGAACCTCTGTAACACACTCAtACtACa CTtATtCCaAGCGCCTGTTTCTTAATCACCATAACCTGCACAT |

**Supplementary Table 11. PCR primers used to generate *J* amplicons.** These are custom primers designed for the first PCR reaction that uses λ genomic DNA as the template. The second PCR step uses standard primers listed in Kelsic *et al.*[13]. Each primer is broken up into three sections. The first, capitalized nucleotides, are the annealing region for the second set of PCR primers. The second, N's of variable length, improve our ability to multiplex amplicons since they cause reading frame shifts so that when PCRs mixed together originating from different primers there will be variability among the clusters on the Illumina flow cell, allowing the machine to more easily distinguish clusters and reduce sequencing errors. The last segment of lowercase letters indicates the chromosome annealing region. The numbers in the primer label (2949.2968 or 3381.3400) indicate which nucleotides the primer anneals to in *J*. Note, *J* is only 3,399 nucleotides long, so 3400 is one nucleotide beyond its reading frame.

| Primer label | Nucleotide sequence |
|---|---|
| J Mage for 2949.2968 6N illum | CCTACACGACGCTCTTCCGATCTNNNNNNgataaacggtacgctgaggg |
| J Mage for 2949.2968 5N illum | CCTACACGACGCTCTTCCGATCTNNNNNgataaacggtacgctgaggg |
| J Mage for 2949.2968 4N illum | CCTACACGACGCTCTTCCGATCTNNNNgataaacggtacgctgaggg |
| J Mage rev 3381.3400 2N illum | GAGTTCAGACGTGTGCTCTTCCGATCTNNctcagaccacgctgatgccc |
| J Mage rev 3381.3400 1N illum | GAGTTCAGACGTGTGCTCTTCCGATCTNctcagaccacgctgatgccc |
| J Mage rev 3381.3400 0N illum | GAGTTCAGACGTGTGCTCTTCCGATCTctcagaccacgctgatgccc |

**Supplementary Table 12.** *P*-values for two-sample t-tests performed to compare fitness effects of the two neutral markers on the corresponding genotype. Genotypes for sites 5, 6 and 9 had data for only one or none replicate population for one of the neutral markers.

| Site | Single-mutation Genotype | P-value | Bonferroni corrected P-value |
|---|---|---|---|
| G1 | 1000000000 | 0.7676 | 1 |
| G2 | 0100000000 | 0.7940 | 1 |
| G3 | 0010000000 | 0.2535 | 1 |
| G4 | 0001000000 | 0.9053 | 1 |
| G5 | 0000100000 | - | - |
| G6 | 0000010000 | - | - |
| G7 | 0000001000 | 0.2988 | 1 |
| G8 | 0000000100 | 0.7333 | 1 |
| G9 | 0000000010 | - | - |
| G10 | 0000000001 | 0.0088 | 0.0616 |

#### References


1 Meyer, J. R. *et al.* Repeatability and contingency in the evolution of a key innovation in phage lambda. *Science* **335**, 428-432, doi:10.1126/science.1214449 (2012).

2 Hendrix, R. W. *Lambda II.* (Cold Spring Harbor Laboratory, 1983).

3 Meyer, J. R. *et al.* Ecological speciation of bacteriophage lambda in allopatry and sympatry. *Science* **354**, 1301-1304, doi:10.1126/science.aai8446 (2016).

4 Boos, W. & Bohm, A. Learning new tricks from an old dog: MalT of the Escherichia coli maltose system is part of a complex regulatory network. *Trends Genet* **16**, 404-409 (2000).

5 Chaudhry, W. N. *et al.* Leaky resistance and the conditions for the existence of lytic bacteriophage. *PLOS Biology* **16**, e2005971, doi:10.1371/journal.pbio.2005971 (2018).

6 Meyer, J. R., Gudelj, I. & Beardmore, R. Biophysical mechanisms that maintain biodiversity through trade-offs. *Nat Commun* **6**, 6278, doi:10.1038/ncomms7278 (2015).

7 Baba, T. *et al.* Construction of Escherichia coli K-12 in-frame, single-gene knockout mutants: the Keio collection. *Mol Syst Biol* **2**, 2006.0008, doi:10.1038/msb4100050 (2006).

8 Datsenko, K. A. & Wanner, B. L. One-step inactivation of chromosomal genes in Escherichia coli K-12 using PCR products. *Proc Natl Acad Sci U S A* **97**, 6640-6645, doi:10.1073/pnas.120163297 (2000).

9 Maddamsetti, R. *et al.* Gain-of-function experiments with bacteriophage lambda uncover residues under diversifying selection in nature. *Evolution* **72**, 2234-2243, doi:10.1111/evo.13586 (2018).
10 Petrie, K. L. *et al.* Destabilizing mutations encode nongenetic variation that drives evolutionary innovation. *Science* **359**, 1542-1545, doi:10.1126/science.aar1954 (2018).
11 Sambrook, J. & Russell, D. W. *Molecular cloning : a laboratory manual*. 3rd edn, (Cold Spring Harbor Laboratory Press, 2001).
12 Wang, H. H. *et al.* Programming cells by multiplex genome engineering and accelerated evolution. *Nature* **460**, 894-898, doi:10.1038/nature08187 (2009).
13 Kelsic, E. D. *et al.* RNA Structural Determinants of Optimal Codons Revealed by MAGE-Seq. *Cell Syst* **3**, 563-571 e566, doi:10.1016/j.cels.2016.11.004 (2016).
14 Benjamini, Y. & Hochberg, Y. Controlling the False Discovery Rate: A Practical and Powerful Approach to Multiple Testing. *Journal of the Royal Statistical Society: Series B (Methodological)* **57**, 289-300, doi:10.1111/j.2517-6161.1995.tb02031.x (1995).
15 R: A Language and Environment for Statistical Computing v. 3.6.1 (R Foundation for Statistical Computing, 2019).
16 Akaike, H. A new look at the statistical model identification. *IEEE Transactions on Automatic Control* **19**, 716-723, doi:10.1109/TAC.1974.1100705 (1974).
17 Drake, J. W. A constant rate of spontaneous mutation in DNA-based microbes. *Proc Natl Acad Sci U S A* **88**, 7160-7164, doi:10.1073/pnas.88.16.7160 (1991).
18 Shao, Y. & Wang, I. N. Bacteriophage adsorption rate and optimal lysis time. *Genetics* **180**, 471-482, doi:10.1534/genetics.108.090100 (2008).
19 Burmeister, A. R., Lenski, R. E. & Meyer, J. R. Host coevolution alters the adaptive landscape of a virus. *Proc Biol Sci* **283**, doi:10.1098/rspb.2016.1528 (2016).
20 Ellis, H. M., Yu, D., DiTizio, T. & Court, D. L. High efficiency mutagenesis, repair, and engineering of chromosomal DNA using single-stranded oligonucleotides. *Proc Natl Acad Sci U S A* **98**, 6742-6746, doi:10.1073/pnas.121164898 (2001).
21 Shuman, H. A. & Silhavy, T. J. The art and design of genetic screens: Escherichia coli. *Nat Rev Genet* **4**, 419-431, doi:10.1038/nrg1087 (2003).
